# Familiarity synchronizes brain rhythms during natural viewing

**DOI:** 10.64898/2026.10.22.760672

**Authors:** Kira N Noad, Daniel Kaiser

## Abstract

Our perception of the world is shaped not only by sensory information, but also by prior experience. However, how familiarity alters the coordination of neural representations across distributed brain networks remains unclear. Rhythmic neural activity provides a potential mechanism for coordinating information across sensory and higher-order brain regions in the light of previous experience. Here, we thus investigated whether familiarity alters the coordination of brain rhythms during naturalistic viewing. Participants who were either familiar or unfamiliar with the television series Game of Thrones viewed clips from the series while EEG or fMRI was recorded. We calculated the intersubject correlations of neural activity across EEG frequency bands and related this rhythmic activity to fMRI BOLD responses. Across participants, all brain rhythms showed reliable intersubject synchrony and related to activity across large-scale brain networks. Familiarity selectively enhanced intersubject synchronisation in beta and gamma rhythms, with these effects reflected in visual and default mode brain regions. Moreover, beta and gamma band activity were modulated by similar aspects of the movie, suggesting coordinated familiarity-related processing. These findings show that familiarity selectively shapes rhythmic neural dynamics across the cortex, providing a mechanism through which prior experience alters the coordination of neural representations during naturalistic perception.

## Introduction

In everyday life, humans must process a continuous stream of dynamic and multi-sensory information to make sense of the world. To face this challenge, the human brain can use familiarity, or prior knowledge, to enhance the processing of incoming sensory information (Jenkins et al., 2011; Kovács, 2020; Manahova et al., 2018; Noad et al., 2024). Prior knowledge generates expectations that influence how sensory information is processed, allowing perception to be guided by both sensory input and knowledge (De Lange et al., 2018).

While we know that familiarity enhances representations across widespread sensory and cognitive brain networks (Epstein et al., 2007; Noad et al., 2024, 2025; Visconti di Oleggio Castello et al., 2021; Wiese et al., 2023), we know much less about how familiarity impacts the orchestration of these representations. Here, we hypothesize that familiarity significantly alters the orchestration of representations through rhythmic activity (Fries, 2015; Varela et al., 2001), providing a mechanism for enhanced, and coordinated, representations of familiar inputs across brain systems. To test this hypothesis, we analyzed EEG and fMRI data recorded during a naturalistic movie watching paradigm and compared groups of participants that were familiar or unfamiliar with the movie.

During naturalistic viewing, time-varying fMRI BOLD activity synchronises between participants (Hasson et al., 2010; Hasson et al., 2004; Hasson, Yang, et al., 2008), and it does so as a function of prior experience, such as familiarity or group membership (Andrews et al., 2019; Noad et al., 2025; van Baar et al., 2021). Familiar movies synchronize activations across a large network of sensory and higher-order regions, likely because of richer conceptual knowledge and stronger predictions compared to unfamiliar movies (Andric et al., 2016; Noad et al., 2024). These findings suggest that familiarity reorganizes neural communication across large-scale neural responses during naturalistic viewing, although the neural mechanisms supporting this altered communication remain unknown.

Rhythmic brain activity provides a candidate mechanism through which familiarity could orchestrate communication across brain networks during naturalistic perception. Large-scale synchronisation of neural rhythms has been linked to information transfer across distributed cortical networks (Fries, 2015; Siegel et al., 2012; Varela et al., 2001), and rhythmic brain activity has been implicated in a wide range of cognitive processes, including sensory processing, memory, and even consciousness (Buzsaki & Draguhn, 2004; Cannon et al., 2014; Engel & Fries, 2016; Keil & Senkowski, 2018; Kucewicz et al., 2024). Some initial EEG studies indeed demonstrate reliable inter-individual synchronization during movie watching, suggesting an involvement of multiple frequency bands (Camenzind et al., 2026; Dmochowski et al., 2012; Ki et al., 2016). Such synchronisation may reflect both shared sensory processing and higher-order processes (Camenzind et al., 2026; Cohen & Parra, 2016; Dini et al., 2023). Different frequency bands may contribute to these processes in different ways: low-frequency rhythms from delta to alpha may reflect changes in sensory inputs at natural timescales, tracking the periodic variability in the environment (Camenzind et al., 2026; Luo et al., 2010). Conversely, synchronization in high-frequency gamma activity may be related to stable feedforward coding and communication of visual feature information (Chen & Farivar, 2020; Kuzovkin et al., 2018; Tallon-Baudry & Bertrand, 1999). At the same time, alpha and beta rhythms have been hypothesized to dynamically coordinate this feedforward-driven synchrony, by routing predictions and coordinating attentional resources (Betti et al., 2021; Fries, 2015; Stecher et al., 2025; Uran et al., 2022). Familiarity with a stimulus may alter the way in which neural rhythms coordinate neural representations: Even when sensory input remains constant for familiar and unfamiliar viewers, familiarity may alter synchronization in both low-frequency alpha/beta and high-frequency gamma dynamics by changing the balance between incoming sensory information and predictions derived from prior knowledge.

Here, we tested this hypothesis by investigating how familiarity shapes rhythmic neural activity across distributed brain networks. We recorded EEG activity while participants who were either familiar or unfamiliar with the television series Game of Thrones viewed clips from the series. We quantified stimulus-driven synchronisation of brain rhythms across individuals and related these dynamics to fMRI responses acquired from an independent group viewing the same stimulus (Noad et al., 2024). We predicted that if brain rhythms support communication between sensory and higher-order systems, familiarity would strengthen both the alignment of brain rhythms across individuals and their relationship to distributed fMRI responses. Across both the familiar and unfamiliar participants, brain rhythms showed reliable intersubject synchronisation and were related to fMRI activity across a large set of brain regions. Crucially, familiarity selectively enhanced intersubject synchrony in beta and gamma rhythms, with these effects localized to both sensory and higher-order regions. Together, these findings suggest that familiarity selectively shapes beta- and gamma-band dynamics across distributed visual and non-visual brain systems, suggesting an altered interplay of sensory information and existing knowledge as naturalistic inputs become familiar.

## Results

We investigated how familiarity shapes rhythmic neural activity across distributed brain networks during naturalistic movie watching. We recorded EEG responses while participants (n = 56, 19-49y) viewed two short movies (movie 1: 12 mins 58s; movie 2: 14 mins 57s) composed of clips from the TV show Game of Thrones (Figure 1A). Participants were either familiar with Game of Thrones (n = 28) or had never seen the show before (n = 28). Participants viewed both movies twice, with the movies presented in the same order in both viewings (movie 1, then movie 2). fMRI responses to movie 1 were acquired from an independent dataset (Noad et al., 2024; https://openneuro.org/datasets/ds004848). Participants in the fMRI group were similarly either familiar (n = 23) or unfamiliar (n = 22) with Game of Thrones.

**Figure 1.**
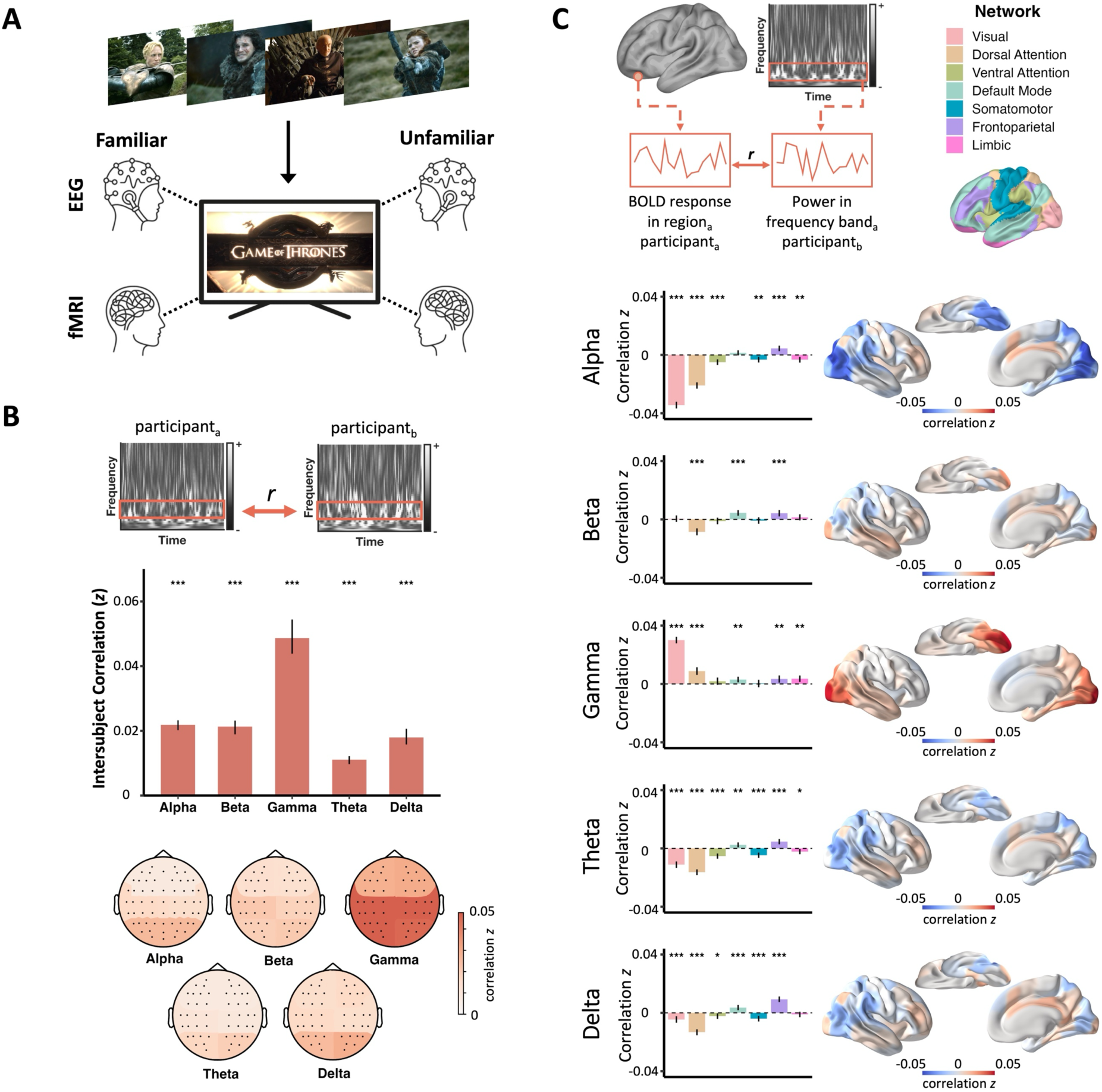
Investigating EEG and fMRI responses during natural viewing. **A**) Participants watched clips from the TV show Game of Thrones while neural responses were measured with EEG or fMRI. Half of the participants were familiar with Game of Thrones, while the other half had never seen the show. **B**) Intersubject correlation (ISC) was used to compare the similarity in neural response between participants. ISC was calculated by correlating time-resolved EEG power in the alpha, beta, gamma, delta, and theta bands across all possible pairs of participants (irrespective of familiarity). ISCs were significantly greater than zero for all rhythms, both when using all EEG electrodes and when splitting them into regions of interest (left/right: posterior, central, frontal). **C**) To obtain a spatial mapping of the rhythmic ISCs, we correlated the time-resolved EEG power with time-resolved fMRI BOLD responses, again across all possible pairs of participants. This analysis was conducted across the brain using the Schaefer et al., (2018) 100-region parcellation and averaged within Yeo (2011) brain networks. The highest correspondence was found in visual regions, with negative correlations for alpha and positive correlations for gamma. All comparisons are corrected with false discovery rate correction (*p* < .001 = ***, *p* < .01 = **, *p* < .05 = *). Error bars reflect 95% confidence intervals from a bootstrapping test.

### Intersubject correlation reveals reliable EEG brain rhythm responses during natural viewing

Before investigating how familiarity alters intersubject synchronization across neural rhythms, we first established whether watching the movies synchronized brain rhythms in the first place. We thus tested the synchronisation of time-varying power in brain rhythms across participants, irrespective of whether they were familiar or unfamiliar with Game of Thrones. Specifically, we performed intersubject correlation (ISC) analysis, which compares the time course of neural responses (here: time-varying rhythmic power) in one participant with the time course of neural responses in another participant, thereby reveal stimulus-driven neural synchronization (Hasson, Landesman, et al., 2008; Hasson et al., 2004).

We first extracted rhythmic responses from the EEG data. We concatenated the first presentation of the two movies into a single timeseries and converted the data into a time-frequency representation, computing rhythmic spectral power using a Hanning taper (frequency range: 1 to 49Hz, in steps of 1Hz; temporal resolution: 100ms; temporal window size: frequency-dependent temporal window corresponding to five cycles for each frequency, ranging from 5s at 1Hz to 102ms at 49Hz). Time-varying power was then averaged for the alpha (8-12 Hz), beta (13-30Hz), gamma (30-49Hz), theta (4-8Hz), and delta (1-4Hz) bands. We then averaged the signal across all electrodes into a single timeseries for each frequency band and participant. To obtains ISCs, we correlated the resulting time courses between all pairs of participants, irrespective of familiarity. Permutation testing was used to assess whether these ISCs were significantly greater than zero (Chen et al., 2016). All brain rhythms showed significant positive ISCs (alpha: *r* = .02, *p* < .001; beta: *r* = .02, *p* < .001; delta: *r* = .02, *p* < .001; gamma: *r* = .05, *p* < .001; theta: *r* = .01, *p* < .001), demonstrating that movie-watching aligns brain rhythms over time across individuals (Figure 1B). To approximate the cortical origin of this synchronization, we repeated the analysis in six regions of interest (ROIs), averaging time-resolved power in electrodes in the left and right posterior, central and frontal areas. The response within each of these ROIs was averaged into a single timeseries and ISCs were again computed for all pairs of participants. ISCs were significantly greater than zero in all regions of interest in all brain rhythms (all *p* < .001; Supplementary Table 1). The strongest ISCs were generally found in posterior regions and the weakest in frontal regions (Figure 1B), in line with fMRI studies showing the greatest neural synchrony in sensory regions (Hasson et al., 2010; Hasson et al., 2004).

Next, we investigated whether similar neural synchronization was observed on the second showing of the movies, too. Similarly to the first showing, ISCs were significantly greater than 0 across all frequency bands (Supplementary Figure 1). However, ISCs were significantly attenuated in the second showing of the movies compared to the first in beta (M_diff_ = .010, *p* < .001), delta (M_diff_ = .006, *p* < .001), and most prominently in gamma (M_diff_ = .043, *p* < .001) rhythms, but showed no significant difference in alpha (M_diff_ = -.003, *p* = .999) and theta (M_diff_ = -.002, *p* = 1) rhythms. This shows rhythmic synchronization was reliable across the two showings, while repetition attenuated the effect, possibly due to a combination of familiarity, adaptation, and fatigue. Since repetition attenuated ISCs, we focused on the first showing for all following analyses.

### Brain rhythms relate to distributed BOLD responses during natural viewing

We have shown that brain rhythms carry stimulus-locked information during natural viewing. However, EEG provides only limited information about the neural sources of these rhythmic responses. Next, we therefore related time-varying EEG power to time-varying fMRI BOLD activity from another dataset, in which participants also watched the first movie shown in the EEG experiment (Noad et al., 2024). This allowed us to relate localized BOLD activity to the global EEG signals in different brain rhythms. As in the EEG study, the fMRI sample comprised 23 participants familiar with Game of Thrones and 22 participants unfamiliar with the show. For the first analysis, we collapsed across all participants, irrespective of familiarity.

We correlated time-varying EEG power (resampled to the fMRI acquisition time; 2s per volume) with the time-varying BOLD responses, separately for each frequency band . We performed this analysis separately for 100 brain parcels (Schaefer et al., 2018), averaging fMRI activity across voxels within each parcel to obtain a single BOLD timeseries. We additionally averaged the correlation between brain rhythms and BOLD response in each parcel within the seven Yeo brain networks (Yeo et al., 2011) to identify patterns at the level of larger-scale functional networks. Figure 1C shows the network-level and whole-brain maps of fMRI-EEG correlations for each brain rhythm, demonstrating significant correlations between time-varying BOLD responses and EEG power in all brain rhythms (for a detailed report, see Supplementary Tables 4 and 5).

The strongest correlations between fMRI and EEG were found in alpha and gamma, in line with previous findings (Goldman et al., 2002; Laufs et al., 2003; Scheeringa et al., 2011; Siegel et al., 2012). Alpha demonstrated an overall pattern of negative correlations with BOLD responses. These negative correlations were strongest within visual regions and became weaker moving from posterior to anterior brain regions. In contrast, gamma showed an overall pattern of positive correlations with BOLD responses, which was especially prominent in visual regions. The spatial pattern largely mirrored that of alpha, with correlations becoming weaker on the posterior to anterior axis. In an additional region-of-interest analysis, negative alpha-to-BOLD and positive gamma-to-BOLD correlations emerged consistently across visual cortex regions (Supplementary Figure 2). Beta rhythms showed generally weak correlations with BOLD response, with no significant correlation in visual regions and no clear overall positive or negative pattern of correlations across the cortex. Delta and theta showed a similar pattern of correlations to each other, characterised by negative correlations with BOLD response in visual and attention regions, and weak positive correlations in default mode and limbic networks and more anterior regions.

### Familiarity enhances intersubject correlation of brain rhythms

Having established that brain rhythms show reliable responses across participants during natural viewing, we next tested whether familiarity effects the synchrony of brain rhythms between participants. We thus split the EEG sample into two groups: participants familiar with Game of Thrones (n = 28), and participants unfamiliar with the show (n = 28).

Before proceeding with this analysis, we first tested whether the familiar and unfamiliar groups really differed in the familiarity with Game of Thrones. To this end, we measured knowledge of the narrative, faces, and scenes from Game of Thrones (Figure 2A). As expected, familiar participants scored significantly higher on the narrative test (*t*(50.7) = 15.7, *p* < .001), on the face recognition test (*t*(28.8) = 22.1, *p* < .001), and on the scene recognition test (*t*(27) = 14.1, *p* < .001).

**Figure 2.**
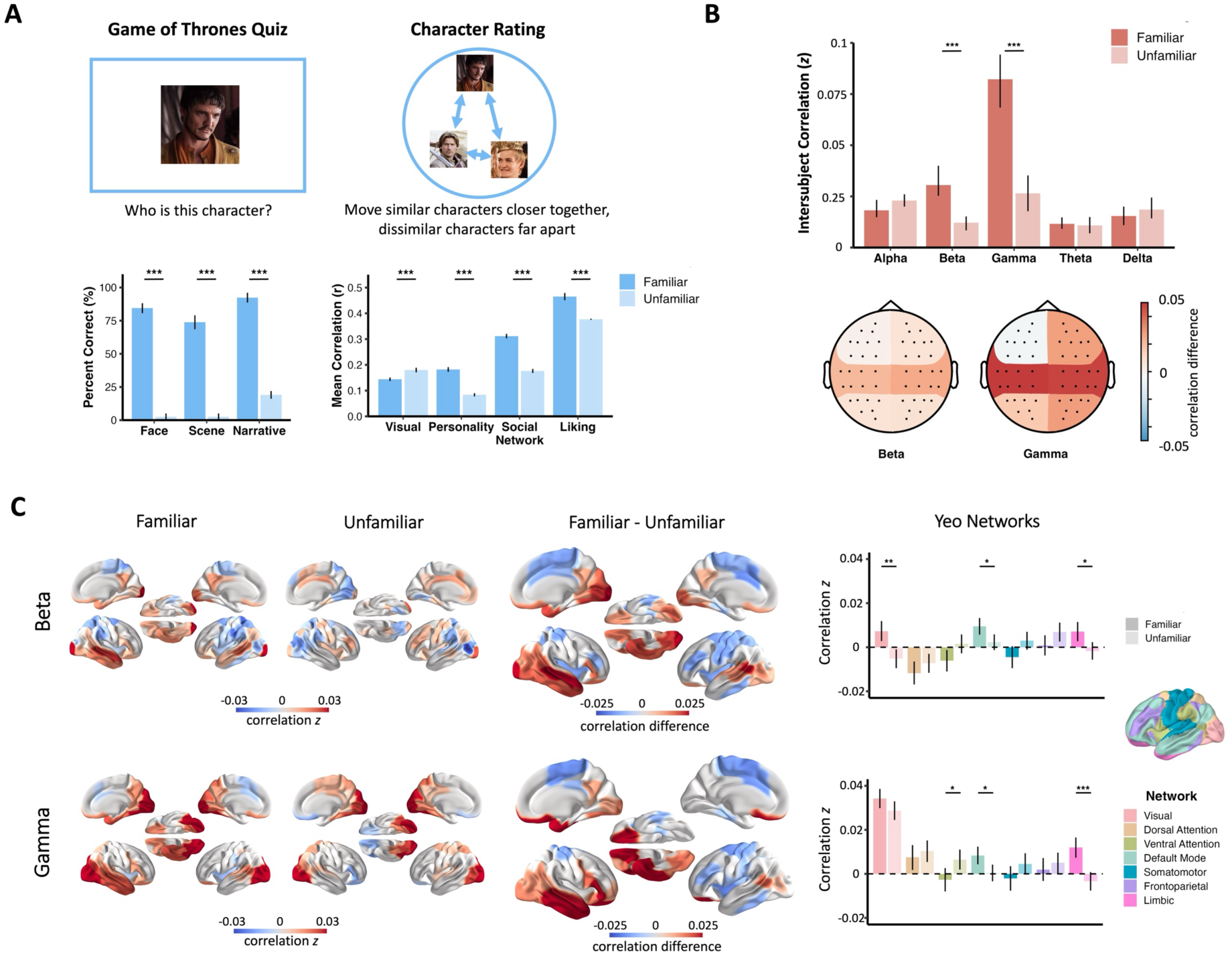
Familiarity alters synchrony in rhythmic brain activity. **A**) Participants completed two behavioural tasks to establish (un)familiarity with Game of Thrones: a general knowledge test, where they answered questions about the narrative, characters and scenes from the show, and a character rating task, where they were asked to sort images of the characters based on how similar they are visually, in their personality, in their social network, and how much they like each character. Familiar participants demonstrated significantly greater knowledge of narrative, faces, and scenes from the show. They were also significantly more aligned in their ratings of characters from the show. **B**) We computed intersubject correlations (ISCs) to compare rhythmic synchronization for the familiar and unfamiliar groups separately. Familiar participants demonstrated significantly greater ISCs in beta- and gamma-rhythms compared to unfamiliar participants, predominantly in central electrodes. **C**) We correlated the time-resolved EEG power in beta and gamma with time-resolved fMRI BOLD responses for the familiar and unfamiliar participants separately. For visualisation, group maps for each group individually are shown in the right hemisphere. The difference between the groups is shown in both hemispheres, with red denoting more positive correlations for familiar participants and blue more positive correlations in unfamiliar participants. Familiar participants demonstrate greater beta- and gamma-rhythm synchrony with BOLD response in visual and higher-order regions, predominantly the visual, default mode and limbic networks. All comparisons are corrected with false discovery rate correction (*p* < .001 = ***, *p* < .010 = **, *p* < .050 = *). Error bars reflect bootstrapped 95% confidence intervals.

We further asked participants to assess how similar characters from the show were visually, in their personality, and in their social network in an arena sorting task (Figure 2A). For each participant, Euclidean distances between character ratings were calculated, and these distances were then correlated between participants to assess the consistency of character representations within each group. Higher correlations indicated greater consistency between participants in how they perceived the characters. Differences in consistency between groups were compared with permutation tests. Unfamiliar participants were significantly more consistent in their visual assessment of the faces (*group difference* = -0.036, *p* = .001), while familiar participants were significantly more consistent in their assessment of personality (*group difference* = 0.098, *p* < .001) and social network similarity (*group difference* = 0.135, *p* < .001). Familiar participants were also more similar than unfamiliar participants in their ratings of how much they liked the characters (*group difference* = 0.260, *p* < .001). These results demonstrate alignment between familiar participants in knowledge about the movie, which is not evident for unfamiliar participants, who seemed to rely more strongly on visual cues.

Having established that the familiar and unfamiliar groups indeed differ in their knowledge of Game of Thrones, we next asked whether familiarity modulates alignment of brain rhythms between participants. To do this, we compared ISCs of brain rhythms separately in the familiar and unfamiliar participants. We thus computed ISCs in each brain rhythm for all combinations of participants in either the familiar or unfamiliar groups. To first check whether synchrony was significant within groups, ISCs were tested against zero in each group using permutation tests. We found that all rhythms showed significant ISCs in both groups across all electrodes, and also in all regions of interest (all *p* < .001) (Supplementary Table 2).

To determine whether familiarity alters synchrony of brain rhythms between participants, we then compared the ISC values between the familiar and unfamiliar groups in each rhythm. Group differences were tested for significance with a permutation test. Whole-brain ISCs were significantly greater for the familiar group compared to the unfamiliar group in beta (*p* < .001) and gamma rhythms (*p* < .001) (Figure 2B), demonstrating that familiarity enhances neural synchrony between subjects. Overall ISC showed no significant difference between groups in alpha (*p* = .064), theta (*p* = .762) and delta (*p* = .351) (Figure 2B). No frequency bands showed greater synchrony for unfamiliar compared to familiar participants. When assessing the topography of the beta and gamma effects across six topographic regions of interest, we observed the greatest differences between the familiar and unfamiliar groups in central electrodes, for both beta and gamma (Figure 2B). Group differences were significant in all regions in beta except the frontal regions and all regions in gamma except the left frontal region (Supplementary Table 3).

### Familiarity alters brain rhythms across cortical networks

Given that familiar participants showed enhanced synchronisation in beta and gamma rhythms, we next used the fMRI data to pinpoint the spatial localization of this enhanced synchronization. Specifically, we tested whether familiar EEG-fMRI correlations (familiar EEG participants correlated with familiar fMRI participants) were significantly different to unfamiliar EEG-fMRI correlations (unfamiliar EEG participants correlated with unfamiliar fMRI participants) using permutation tests. This difference was again calculated on the whole brain (100 parcels, Schaefer et al., 2018), and network level (7 networks, Yeo et al., 2011).

Familiarity significantly altered the relationship between time-varying beta and gamma power and regional BOLD responses. Figure 2C shows BOLD-EEG correlations split by familiarity. Familiar participants showed substantial BOLD-EEG correlations for both beta and gamma rhythms, most prominently characterised by positive correlations in visual and temporal regions (Supplementary Table 6). In contrast, unfamiliar participants showed small, or even negative correlations in these regions (Supplementary Table 7). Comparing EEG-fMRI ISCs across groups revealed significant group differences throughout the ventral visual stream and into anterior brain regions, particularly in the right hemisphere. Strikingly, beta rhythms were only marginally related to BOLD responses when pooling across all participants in our earlier analysis (Figure 1C). Yet, splitting the group into familiar and unfamiliar participants revealed near opposite effects.

To investigate whether these familiarity effects were organised into large-scale brain networks, we averaged the correlation values between the brain rhythms and BOLD response of the Schaefer parcels within each of the 7 Yeo networks. Familiar participants showed significantly greater positive correlations than unfamiliar participants between gamma and BOLD responses in the default mode and limbic networks (default mode: observed difference *r* = .008, *p* = .021; limbic: observed difference *r* = .015, *p* < .001) (Figure 2C, Supplementary Tables 8, 9, and 10). Familiar participants also showed significantly greater correlations than unfamiliar participants between beta power and BOLD responses in the visual, default mode, and limbic networks (visual: observed difference *r* = .012 , p = .001; default mode: observed difference *r* = .002, *p* = .023; limbic: observed difference *r* = .009, *p* = .017). The other frequency bands (alpha, delta, theta) showed comparatively small or no group differences (Supplementary Figure 3, Supplementary Table 10). Together, these results suggest that beta and gamma rhythms are involved in coordinating familiarity-related processing across large-scale brain networks, from sensory to higher-order regions.

### Synchronisation of ISCs across different brain rhythms

So far, we have shown that brain rhythms synchronise across viewers during natural viewing, and that familiarity enhances the synchronisation in beta and gamma rhythms. But do these rhythms respond to the input in similar ways, or do they respond to different aspects of the stimulus? For example, ISCs at timepoint 1 may be high for brain rhythm A and low for brain rhythm B, while ISCs at timepoint 2 may be low for brain rhythm A and high for brain rhythm B. This would suggest different aspects of the movie drive alignment between participants for different rhythms. In contrast, it may be that different brain rhythms show a similar temporal distribution of ISCs. For example, ISCs at timepoint 1 may be high for both brain rhythm A and B, while ISCs at timepoint 2 may be low for both brain rhythm A and B. This would suggest similar aspects of the movie driving alignment between participants in different rhythms.

To test for such inter-rhythm synchrony, we calculated spatial ISCs by correlating the pattern of responses across all electrodes in one brain rhythm at one timepoint in one participant, with the pattern of response in the same brain rhythm at the same timepoint in another participant (Figure 3A left). This was repeated for all combinations of participants at all timepoints, yielding an ISC time course over the duration of the movie for each brain rhythm, which can then be compared across rhythms (Figure 3A right). This spatial ISC provides insights into similarities in the multielectrode pattern of activity at each timepoint, rather than simply asking whether the overall signal fluctuates similarly over time. Thus, two rhythms may show high spatial ISC at the same timepoints even when their absolute power or spatial patterns differ, provided that each rhythm shows a consistent pattern of activity across viewers. Correlating these spatial-ISC time courses therefore allows us to determine whether different brain rhythms show synchronisation between viewers at similar or different moments during the movie. In contrast, a simple correlation of power over time may not be sensitive to these effects (Supplementary Figure 4).

**Figure 3.**
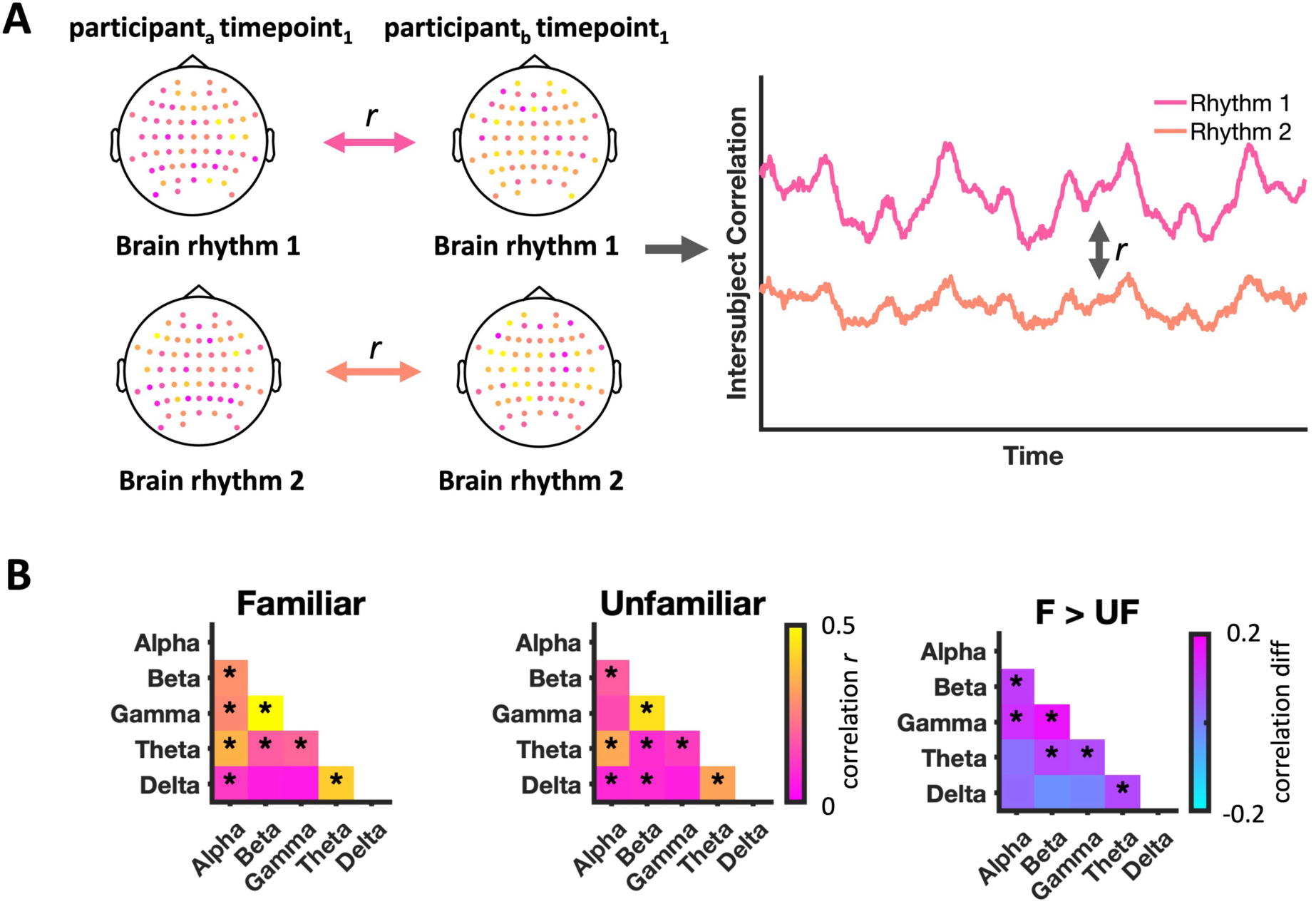
Synchronisation of ISCs across brain rhythms. **A**) Spatial intersubject correlation was calculated by correlating the pattern across electrodes at one timepoint in one participant with the pattern across electrodes at the same timepoint in another participant. This was repeated for all participants, at all timepoints, in each brain rhythm. The correlations across all subjects were averaged at each timepoint, yielding a time course of spatial ISCs for each rhythm. To assess whether brain rhythms are synchronised in their temporal dynamics, the ISC time courses were correlated across rhythms. **B)** Significant positive correlations between ISCs across brain rhythms were found for both familiar and unfamiliar participants. Correlations between beta and gamma rhythms, as well as between beta, gamma, and other rhythms, were significantly greater for familiar than unfamiliar participants. All comparisons are corrected with false discovery rate correction (*p* < .001 = *).

We were interested in two possible outcomes: 1) whether ISCs in brain rhythms showed synchronised temporal dynamics, with increased and decreased strength at the same timepoints in the movie, or 2) whether ISCs showed desynchronised temporal dynamics between the brain rhythms, with different aspects of the movie eliciting ISCs for different rhythms. To assess this, we correlated the average ISC over time in each pair of brain rhythms for familiar and unfamiliar participants. Significance within each group and group differences were tested with a permutation test. The inter-rhythm ISCs were significantly positively correlated in both familiar and unfamiliar groups, demonstrating synchronisation in temporal dynamics (Figure 3B, left, middle; Supplementary Table 12). No band pairings showed significant negative correlations or entirely desynchronised temporal dynamics.

We next tested whether familiarity altered the synchronisation of ISCs across brain rhythms. ISC correlations between beta and gamma were significantly greater for familiar compared to unfamiliar participants (correlation difference = 0.17, *p* <.001) (Figure 3B, right), demonstrating that familiarity effects are related to a joint temporal coupling of these frequency bands. Correlations including beta or gamma and other rhythms were also greater for familiar participants (beta-theta: correlation difference = 0.09, *p* <.001; beta-alpha: correlation difference = 0.10, *p* <.001; gamma-theta: correlation difference = 0.0.8, *p* <.001; gamma-alpha: correlation difference = 0.12, *p* <.001; theta-delta: correlation difference = 0.08, *p* <.001). No band pairings showed greater correlations for unfamiliar compared to familiar participants (Supplementary Table 12).

## Discussion

Perception engages multiple brain rhythms and distributed cortical systems, and familiarity is known to shape neural responses to naturalistic stimuli. However, the neural mechanisms coordinating the enhanced synchronization to familiar stimuli have remained unclear. Rhythmic neural activity provides a potential mechanism for orchestrating aligned brain responses, facilitating communication between distributed brain systems. Here, we put this idea to the test, combining intersubject synchronisation of EEG brain rhythms with fMRI BOLD activity during naturalistic movie viewing. We show that naturalistic perception is supported by coordinated neural dynamics that are shared across individuals and systematically linked to distributed cortical networks. Although stimulus-locked responses were evident across all brain rhythms and a large network of brain regions, familiarity selectively enhanced beta- and gamma-band synchronisation and strengthened their relationship to BOLD responses in sensory and higher-order cortical regions. These findings suggest that familiarity reorganises the coordination of neural responses across distributed brain networks.

### Naturalistic inputs synchronize rhythmic neural activity

The finding that stimulus-locked responses were present across all brain rhythms suggests that naturalistic perception recruits multiple rhythmic channels of neural communication. This extends previous work showing that visual information is coded across low- and high-frequency brain activity (Clarke et al., 2018; Kaiser, 2022; Reddy et al., 2021; Schyns et al., 2011) by demonstrating that these rhythmic dynamics are synchronised across individuals during natural viewing. The engagement of multiple brain rhythms likely reflects simultaneous recruitment of multiple levels of processing during movie watching, from shared sensory responses to higher-order processes involved in interpreting narrative. Consistent with this interpretation, sensory information such as visual and audio-visual processing is known to engage multiple frequency bands (Luo et al., 2010; Moinnereau et al., 2020). Likewise, higher-order cognitive functions are known to synchronise EEG signals between individuals (Cohen & Parra, 2016; Dini et al., 2023) and to engage theta and gamma rhythms (Chang et al., 2015; Dmochowski et al., 2012; Wang et al., 2016). The observed ISCs therefore likely reflect coordinated neural communication supporting both shared perceptual processing and shared interpretation of the narrative. This parallels findings from fMRI studies, where the strongest ISCs are often observed in sensory regions (Hasson et al., 2010; Hasson et al., 2004), whereas manipulations of narrative such as familiarity produce the largest changes in higher-order cortical regions (Noad et al., 2024; Regev et al., 2013; Yeshurun et al., 2017; Zadbood et al., 2017).

### Rhythmic synchronization is linked to widespread regional activity

Correlating time-varying rhythmic activity with time-varying fMRI BOLD, we observed robust covariations of EEG power and BOLD responses in alpha and gamma rhythms, with alpha showing negative correlations and gamma showing positive correlations. These effects were strongest in visual brain regions and were observed bilaterally. These results align with previous findings demonstrating that alpha typically shows a negative correlation with BOLD, while gamma typically shows a positive correlation (Goldman et al., 2002; Laufs et al., 2006; Muthukumaraswamy & Singh, 2008; Xavier et al., 2025) and extends these findings to naturalistic, stimulus-driven conditions using a between-subjects approach. By linking brain rhythms and fMRI responses across participants viewing the same stimulus, our results demonstrate that these relationships are reliably driven by the stimulus rather than idiosyncratic within-subject fluctuations. The convergence between rhythmic activity and distributed BOLD responses suggests that electrophysiological and haemodynamic measures capture complementary aspects of the same large-scale changes unfolding during naturalistic perception.

### Familiarity enhances synchronization in beta and gamma rhythms

Familiarity selectively enhanced neural synchrony. While all brain rhythms were synchronised in both familiar and unfamiliar participants, only beta- and gamma-bands showed greater intersubject synchrony for familiar viewers compared to unfamiliar ones. Synchronisation in the remaining frequency bands did not differ between groups. This pattern is consistent with previous fMRI work using the same stimulus, which showed that familiarity synchronises neural responses in higher-order cortical regions between subjects (Noad et al., 2024). These findings indicate that the effects of familiarity observed at the level of distributed BOLD responses may be supported by frequency-specific coordination of neural dynamics. Familiar stimuli may engage richer conceptual and predictive representations (Bar et al., 2006; De Lange et al., 2018) and recruit stronger interplay between perceptual and conceptual systems (Bar, 2004; Brookshire et al., 2021; Gobbini & Haxby, 2007). Our results suggest that these interactions are supported by coordinated oscillatory dynamics, providing a neural mechanism through which familiarity facilitates communication between distributed brain regions during naturalistic perception.

The selective modulation of beta and gamma suggests that familiarity does not enhance neural communication indiscriminately, but instead acts on brain rhythms that have previously been proposed to coordinate communication between different levels of the cortical hierarchy. Gamma-band activity has been associated with feedforward signalling from lower to higher-order cortical regions (Bartoli et al., 2019; Bastos et al., 2015; Van Kerkoerle et al., 2014), whereas beta-band activity has been linked to feedback communication conveying prior knowledge and predictions (Bastos et al., 2015; Betti et al., 2021; Mejias et al., 2016; Michalareas et al., 2016). Within this framework, enhanced gamma synchronisation in familiar participants may reflect more reliable feedforward communication of sensory information, leading to more consistent stimulus-driven responses across individuals. In contrast, increased beta synchronisation may reflect stronger feedback communication from higher-order systems, consistent with familiarity enabling more accurate predictive signals. This interpretation is further supported by the EEG-fMRI coupling results, which demonstrated that beta and gamma covaried with BOLD response in both sensory and higher-order regions in familiar participants, consistent with their involvement in processing across multiple levels of the cortical hierarchy.

The EEG-fMRI coupling results show familiarity effects of gamma and beta in the visual cortex, fitting with previous work which indicates that familiarity can influence visual processing from early stages of the cortical hierarchy (Friese et al., 2012; Grill-Spector et al., 2006; Manahova et al., 2018). Of particular interest, beta-band activity in visual cortex did not show an overall familiarity-related increase in synchronisation, but instead showed opposing patterns across the familiar and unfamiliar groups: Beta was positively correlated with BOLD in the familiar group but negatively in the unfamiliar group. Given the proposed role of beta in feedback communication within the visual hierarchy (Bastos et al., 2015; Michalareas et al., 2016), this pattern tentatively indicates that familiarity tips the balance of top-down and bottom-up flows in favour of top-down predictions enabled by prior knowledge.

Familiarity-related effects also extended to higher-order regions, including default mode and limbic areas. These effects may reflect enhanced interactions between sensory and higher-order systems, through which incoming visual information can be integrated with prior conceptual, episodic and affective knowledge. This interpretation is consistent with our previous fMRI findings using the same stimulus, where familiarity increased functional connectivity between visual regions and non-visual regions associated with semantic, episodic and affective processing (Noad et al., 2024). Naturalistic viewing studies further show that default-mode and limbic regions integrate narrative meaning, memory and emotional information during movie viewing (Chen et al., 2017; Yang et al., 2023; Yeshurun et al., 2017). Thus, familiarity-related enhancement of visual and non-visual regions may provide a mechanism through which prior experience enriches the interpretation of ongoing sensory input. Future work using approaches capable of resolving the direction and content of these interactions will be important for testing this possibility. Together, our findings suggest that familiarity strengthens communication across the cortical hierarchy and may enhance both the bottom-up transmission of sensory information and the top-down integration of prior knowledge.

If beta and gamma rhythms indeed support efficient convergence between bottom-up and top-down channels of communication within the cortical hierarchy, they would further be expected to exhibit coordinated temporal dynamics. Consistent with this prediction, beta and gamma showed highly aligned fluctuations in ISCs over time, particularly in familiar participants, indicating that beta- and gamma-band synchronisation increased and decreased at similar moments in the movie. This is also in line with beta and gamma rhythms exhibiting comparable patterns of coupling with distributed BOLD responses. These findings suggest that beta and gamma may operate as coordinated components linking sensory processing with higher-order cortical representations during naturalistic perception.

Synchronization in other frequency bands was not altered by familiarity, suggesting that synchronization beyond the beta and gamma bands was primarily driven by the stimulus statistics rather than by familiarity. This is notable given that alpha, like beta, has been implicated in content-related processing and predictive signalling (Fries, 2015; Stecher et al., 2025). However, alpha and beta may not support identical forms of feedback. Work examining the laminar organisation of cortical communication suggests distinct relationships between alpha- and beta-band activity and cortical connectivity, with beta activity implicated in inter-regional feedback pathways (Bastos et al., 2020; Scheeringa et al., 2023). Further work suggests that alpha and beta may convey different forms of top-down information (Di Dona & Ronconi, 2023; Sedley et al., 2016). Thus, although both rhythms have been linked to top-down processing, their functional and anatomical distinctions need further investigation, which may ultimately yield insights into why familiarity selectively modulates beta-band, rather than alpha-band activity.

### Brain rhythms as indicators of interpersonal alignment

More broadly, the present findings highlight the potential of brain rhythms as markers of interpersonal alignment. Intersubject synchronisation is thought to reflect the extent to which individuals respond similarly to a shared stimulus (Finn et al., 2020; Nastase et al., 2019). While this synchronisation predominantly reflects sensory processing, previous work has shown that shared neural dynamics can reflect shared interpersonal factors, such as subjective experience, preferences, understanding, and even personality (Finn & Bandettini, 2021; Liu et al., 2019; Nguyen et al., 2019; van Baar et al., 2021). The present findings suggest that rhythmic brain activity may also provide a measure of interpersonal alignment that is shaped not only by shared sensory input, but by the shared experience of the individuals. Here, we demonstrate that familiarity shapes interpersonal alignment; however, familiarity represents just one aspect of prior experience. Future research could examine how differences in knowledge, preferences, transient states, and stable individual traits influence rhythmic alignment. Comparing the similarity of rhythmic responses across individuals could provide a way to test hypotheses about shared subjective experiences and preferences, and ultimately help explain why people sometimes experience the world in similar ways and sometimes in markedly different ways.

## Conclusion

By linking stimulus-driven rhythmic responses to distributed fMRI responses, our work demonstrates how rhythmic brain activity relates to large-scale brain networks, and suggests that familiarity may alter communication across the brain through selective modulation of rhythmic activity. Familiarity specifically reshapes the coordination of neural responses through synchronized and coupled beta- and gamma-band dynamics. More broadly, these findings suggest that rhythmic neural responses are a mechanism through which prior experience shapes distributed neural responses to shared sensory input.

## Methods

### Participants

Fifty-six participants (median age: 26, age range: 19-49, 18 male) took part in the EEG study. All participants were neurologically healthy by self-report, had normal or corrected-to-normal vision, and were English speakers. Twenty-eight participants had previously watched the TV series Game of Thrones (GoT). The remaining 28 participants had not watched GoT. The sample size was determined a priori based on prior neuroimaging studies recruiting movie watching and similar analysis techniques (Andrews et al., 2019; Hasson, Yang, et al., 2008; Noad et al., 2024, 2025). Written informed consent was obtained for all participants, and the study was approved by the general ethical committee of Justus Liebig University Giessen.

### Stimuli

Participants viewed two movies constructed from audio-visual segments from Seasons 2, 3 and 4 of GoT. The first movie had a length of 12 min 58s (the same movie used in Noad, Watson & Andrews, 2024), featuring 10 distinct scenes ranging in length from 50 to 117s. The second movie had a length of 14 min 57s, featuring 11 distinct scenes ranging in length from 40 to 226s.

Static face images of 9 characters featured in the videos (10 images per character, totalling 90 unique images) were also shown to participants in a passive viewing task. Each image appeared 20 times, resulting in 1800 static image trials per participant. These data are not reported in this paper, as they were collected under a separate research question targeting identity-specific neural representations.

### Procedure

We recorded 64-channel EEG while participants freely viewed the GoT videos on a 32-inch Gigabyte AORUS FI32G monitor. Participants were seated 60cm from the screen and used a chinrest to maintain consistent viewing distance and minimise head movement. Participants watched the first GoT movie, and then the second GoT movie. Both videos subtended approximately 16 x 9 degrees of visual angle on a black background. Following this, participants viewed static face images of characters featured in the videos. Participants then viewed the two GoT videos for the second time, under identical conditions to the first viewing. All stimuli were presented using PsychoPy (Peirce et al., 2019).

All participants performed a behavioural test prior to the EEG session to determine their familiarity with GoT. Understanding of the narrative of the show was assessed in a set of 14, 4-alternative choice questions. Participants also viewed images of faces and scenes from GoT and were asked to name the face/scene or provide identifying information. All tests were self-paced.

Finally, participants performed a behavioural sorting task. Images of the faces of 17 characters from the movie were presented in an arena task, and participants were asked to sort the images so that characters who were more similar were placed more closely together, and characters who were more dissimilar were placed further apart. Participants repeated this three times, sorting the faces based on how visually similar they are, on how similar the characters are in their personality, and based on how similar the characters’ social networks are. Participants were also asked to rate how much they liked each of these characters on a scale of 0-1 (0 being strongly dislike, 1 being strongly like).

### EEG Acquisition and Preprocessing

EEG was recorded using a 64-channel ActiCAP system with an ActiCHamp amplifier (Brain Products GmbH, Gilching, Germany) at a 1000Hz sampling rate. Electrodes were positioned according to the international 10-10 system. Preprocessing was performed using FieldTrip (Oostenveld et al., 2011). Data were epoched from -1 seconds to the end of the movie. Band-stop filtering at 48-52Hz was applied to remove line noise. Data were then re-referenced, downsampled to 200Hz and baseline corrected using the pre-stimulus interval. Noisy channels were identified by visual inspection and removed. Blink-related artifacts were removed using independent component analysis (ICA).

Frequency decomposition was conducted for each GoT video for each participant. For each video presentation, the whole movie period was transformed into the time-frequency domain in each channel using FieldTrip. Power estimates across frequencies were obtained using the Hanning taper from 1 to 49Hz in 1Hz steps. An adaptive sliding time window was used, resulting in longer windows for lower frequencies, and shorter windows for higher frequencies (five cycles for each frequency, ranging from 5s at 1Hz to 102ms at 49Hz). Power was estimated every 100ms across the entire movie. For band-limited analyses, power estimates were averaged across frequency ranges, corresponding to 1-4Hz for delta, 4-8Hz for theta, 8-12Hz for alpha, 13-30Hz for beta and 30-49Hz for gamma. Channels were averaged into 6 regions-of-interest (left/right: frontal, central and posterior), and into an overall signal across all channels.

### Intersubject correlation of brain rhythm power

Intersubject correlations (ISCs) were calculated on power over time in each brain rhythm. This was calculated for each movie presentation separately. To do this, the time-varying power in one brain rhythm in one participant was correlated with the time-varying power in the same brain rhythm in another participant (Pearson’s *r*) across the whole movie. This was repeated for all combinations of participants, and for each brain rhythm. A Fisher’s *z* transform was applied to the correlations prior to statistical analysis.

First, we tested whether brain rhythm ISCs were significantly greater than 0. A sign-flip permutation test was conducted to assess significance. This test assumes that positive and negative values are equally likely under the null hypothesis and therefore creates a null distribution of correlation values by randomly assigning positive and negative signs (5000 permutations used). The true median ISC value was tested against the median permuted ISC value to test significance. Median rather than mean ISC is used as it is more appropriate for non-parametric tests (Chen et al., 2016; Nastase et al., 2019). ISC was calculated on the overall power averaged across all channels, and for the 6 regions of interest. Permutation tests were calculated for the overall power averaged across all channels for each brain rhythm separately, and for each ROI in each brain rhythm (with Bonferroni-Holm correction across ROIs within a brain rhythm for multiple comparisons). 95% confidence intervals were calculated using bootstrapping (5000 iterations with replacement).

To then compare whether ISCs differ between the familiar and unfamiliar groups in any frequency bands, group assignment permutation tests were conducted. This test randomly assigns group membership at each permutation, under the assumption that there will be no difference between groups under the null hypothesis. The true observed group difference median ISCs (the median ISC of the familiar group – the median ISC of the unfamiliar group) is then compared to the permuted group difference ISCs (5000 permutations) to test whether there is a significant difference between groups. Group difference permutation tests were calculated for the overall power averaged across all channels for each brain rhythm separately, and for each ROI in each brain rhythm (with Bonferroni-Holm correction across ROIs within a brain rhythm). 95% confidence intervals were calculated using bootstrapping (5000 iterations with replacement).

### fMRI Dataset and Preprocessing

We next sought to compare time-varying power in each brain rhythm to fMRI BOLD response over time. We used the publicly available Game of Thrones MRI dataset (https://openneuro.org/datasets/ds004848), described previously in Noad, Watson & Andrews (2024). This dataset includes 45 neurologically healthy participants with normal or corrected-to-normal vision (15 male, 30 female, age range = 18-32, median age = 19). Twenty-two of the participants had watched GoT, and 23 participants had not watched GoT. Participants passively viewed a video comprising of clips from Game of Thrones. Videos were presented at the full resolution of the screen (1920 × 1080 pixels, subtending approximately 38.7° × 22.3° visual angle). The GoT video presented in MRI is the same as the first video presented to the EEG participants.

MRI data were collected at the York Neuroimaging Centre on a 3T Siemens Magnetom Prisma scanner using a 64-channel head coil. Functional data were acquired from 60 axial slices using a multiband EPI sequence (TR = 2 s, TE = 30 ms, FOV = 240 × 240 mm, matrix size = 80 × 80, 3×3×3mm voxels, flip angle = 80°, anterior-posterior phase encoding direction, multiband acceleration factor = 2). Additional field-map images were acquired in the same plane as the functional images (TR = 554 ms, short TE = 4.9 ms, long TE = 7.38 ms, flip angle = 60°, other parameters as per functional images). T1-weighted anatomical images were acquired from 176 sagittal slices (TR = 2.3 s, TE = 2.26 ms, FOV = 256 × 256 mm, matrix size = 256 × 256, 1x1x1mm voxels, flip angle = 8°).

MRI data were pre-processed using FSL (Jenkinson et al., 2012). Motion correction using MCFLIRT (Jenkinson et al., 2002), temporal high-pass filtering (Gaussian-weighted least squares straight line fittings, sigma = 50s) and slice timing correction were applied. Spatial smoothing (Gaussian) was applied at 6mm full width at half maximum, and non-brain material was removed using BET (Smith, 2002). Functional images were then co-registered to the T1 anatomical images via boundary-based registration (Greve & Fischl, 2009). Field-maps were used to additionally apply B0 unwarping to the functional images. Anatomical images were then registered to the MNI152 brain via a nonlinear registration using FNIRT (Andersson et al., 2010).

### Relating time-varying EEG power to BOLD response

Most studies rely on within-subject correlations between EEG power and BOLD response, which are sensitive to noise and exhibit substantial inter-individual variability (Meyer et al., 2013). Between-subject approaches offer a complementary framework by isolating neural dynamics that are shared across individuals. Leveraging a between-subjects approach, we correlated the BOLD response over time in each region of the brain with the power in each brain rhythm over time for each pair of participants (Figure 1C). To compare BOLD response to rhythmic EEG power, we first downsampled the EEG data to match the temporal resolution of the fMRI data. The power in each brain rhythm was averaged within consecutive 2 second bins to align with the 2 second TR in the fMRI data, leading to a total of 389 “timepoints” in both the EEG and fMRI data. Since fMRI relies on the slow haemodynamic response, EEG responses were temporally shifted by 6 seconds. BOLD response was partitioned based on the Schaefer 100 brain parcellations (Schaefer et al., 2018), and the normalised timeseries was averaged across all voxels within each parcel to produce one BOLD timeseries per participant in each parcel.

For each pair of EEG and fMRI participants, the power in each brain rhythm over time was correlated (Pearson’s *r*) with the BOLD response over time within a region of the brain. Different participants took part in the fMRI and EEG studies, making all pairings between-subject. The correlation was repeated for all pairs of participants, for all brain rhythms and all regions of the brain. Correlations were converted to Fisher’s *z* prior to statistical analysis.

First, we sought to test whether time-varying power in any brain rhythms significantly correlated with BOLD response. In each brain rhythm, we tested whether correlations were significantly different from zero using a sign-flip permutation test. This test assumes that positive and negative values are equally likely under the null distribution, and randomly flips the sign of the values at each permutation to build a null distribution. We did this on a network level using Yeo’s 7 networks (Yeo et al., 2011), and for each region in the Schaefer 100 parcels. At the network level, we averaged correlations across participant pairs to produce one correlation per region within the network. At the region level, we used all pairwise correlations to test whether values were significantly different to zero. False discovery rate correction was applied to correct for multiple comparisons within each brain rhythm. 95% confidence intervals were calculated using bootstrapping (5000 iterations with replacement).

Next, we tested whether correlations between time-varying power in EEG brain rhythms and BOLD response differed based on familiarity. Here, we tested whether familiar pair correlations (i.e. familiar EEG participant with familiar fMRI participant) were significantly different to unfamiliar pair correlations (i.e. unfamiliar EEG participant with unfamiliar fMRI participant) using a group assignment permutation test. This test assumes there to be no difference between the groups under the null hypothesis, and randomly assigns group membership at each permutation to create a null distribution of group differences. This was performed on a network level (averaging correlations across pairs of participants to produce one correlation value per region per group within the network) and per region (comparing all pairwise correlations in one group to all pairwise correlations of the other group). False discovery rate correction was applied to correct for multiple comparisons within each brain rhythm. 95% confidence intervals were calculated using bootstrapping (5000 iterations with replacement).

### Synchronisation of ISCs across different brain rhythms

To explore how intersubject correlations varied over time, we calculated spatial ISCs for each brain rhythm. To do this, the pattern of response across electrodes in each brain rhythm at one timepoint in one participant was correlated with the response across electrodes at the same timepoint in the same brain rhythm in another participant. This was repeated for all pairs of participants, at all timepoints, and in all brain rhythms, across the first repetition of the movies. Correlation values were Fisher’s *z* transformed and averaged between participant pairs at each timepoint, resulting in a single intersubject correlation over time. Since preprocessing involved removing noisy channels, not all channels were present in all subjects. Therefore, the intersection of shared channels for each individual pair was used to calculate spatial ISCs.

We sought to understand whether ISCs fluctuated in similar or dissimilar ways across the brain rhythms. We tested two potential outcomes: 1) different brain rhythms may show synchronised temporal dynamics over time, with peaks and troughs in ISCs at similar timepoints in the movie, or 2) brain rhythms may show desynchronised temporal dynamics, with different aspects of the movie driving alignment between participants for different rhythms. To determine synchrony between rhythms, we correlated the ISC over time between each rhythm (i.e. Beta ISC over time with Gamma ISC over time). This was calculated separately for each participant group. Statistical significance of between brain rhythm correlations was assessed using a permutation test (1,000 permutations). For each brain rhythm pair (e.g. beta-gamma), the observed Spearman correlation between ISC time courses was compared to a null distribution generated by circularly shifting one time series randomly and recomputing the correlation. P-values were computed as the proportion of permuted correlations with absolute values greater than or equal to the observed correlation.

Finally, we tested whether familiar participants showed more temporal alignment between rhythms than unfamiliar participants. To calculate whether correlations between brain rhythms were different between the groups, group assignment permutation tests were conducted. For each group, ISC time courses were first averaged across participant pairs within that group, and Spearman correlations were computed between all pairs of frequency bands to obtain rhythm-to-rhythm correlation matrices. To generate a null distribution, the group labels assigned to participant pairs were randomly permuted (1,000 permutations) while keeping the total number of pairs in each group constant. For each permutation, the group difference in the rhythm-to-rhythm correlations was stored. Statistical significance was assessed by comparing the observed correlation difference to the permuted null distribution.

As a comparison, the simple correlation of power over time between each rhythm was also calculated within a participant. The overall power across electrodes was used, and correlated across timepoints between rhythms. Statistical significance of between rhythm correlations was assessed using a permutation test (1,000 permutations). For each brain rhythm pair (e.g. beta-gamma), the observed Spearman correlation between ISC time courses was compared to a null distribution generated by circularly shifting one time series randomly and recomputing the correlation. P-values were computed as the proportion of permuted correlations with absolute values greater than or equal to the observed correlation. This was computed for each group (familiar, unfamiliar) to demonstrate whether correlations were significantly greater than the null. We next tested whether familiar participants showed more alignment between rhythms than unfamiliar participants. To calculate whether correlations between brain rhythms were different between the groups, group assignment permutation tests were conducted. For each group, the time courses were first averaged across participant pairs within that group, and Spearman correlations were computed between all pairs of frequency bands to obtain rhythm-to-rhythm correlation matrices. To generate a null distribution, the group labels assigned to participant pairs were randomly permuted (1,000 permutations) while keeping the total number of pairs in each group constant. For each permutation, the group difference in the rhythm-to-rhythm correlations was stored. Statistical significance was assessed by comparing the observed correlation difference to the permuted null distribution.

## Data and code availability

Data (https://doi.org/10.5281/zenodo.23396662) and code (https://doi.org/10.5281/zenodo.23393613) are freely available on Zenodo.

## Acknowledgements

This work was supported by the Deutsche Forschungsgemeinschaft (DFG), grants KA4683/5-1 (project no. 518483074), KA4683/7-1 (project no. 548389777), and under Germany’s Excellence Strategy (EXC 3066/1, “The Adaptive Mind”, project no. 533717223). It was further supported by a European Research Council (ERC) Starting Grant (PEP, ERC-2022-STG 101076057). Views and opinions expressed are those of the authors only and do not necessarily reflect those of the funders. Neither the funders nor the granting authority can be held responsible for them. We would like to thank Tanja John for help with EEG data collection, and Tim Andrews and David Watson for the previous collection of fMRI data.

