## Supplementary Tables and Figures for "Familiarity synchronizes brain rhythms during natural viewing"

**Supplementary Table 1.** Intersubject correlation values for all subjects in the frequency bands, split by electrode region. All correlations were significantly greater than 0, tested with a sign-flip permutation test and Bonferroni-Holm corrected within a band.

| Hemisphere | Region | Alpha | Beta | Delta | Gamma | Theta |
|---|---|---|---|---|---|---|
| L | Posterior | .026 | .020 | .024 | .045 | .017 |
| R | | .027 | .020 | .028 | .040 | .020 |
| L | Central | .016 | .022 | .016 | .060 | .010 |
| R | | .014 | .018 | .013 | .045 | .010 |
| L | Frontal | .008 | .012 | .015 | .025 | .010 |
| R | | .008 | .012 | .015 | .020 | .010 |

**Supplementary Table 2.** Intersubject correlation values for familiar (F) and unfamiliar (UF) subjects in the frequency bands, split by electrode region. All correlations were significantly greater than 0, tested with a sign-flip permutation test and Bonferroni-Holm corrected within a band.

| | | Alpha | | Beta | | Delta | | Gamma | | Theta | |
|---|---|---|---|---|---|---|---|---|---|---|---|
| Hemisphere | Region | F | UF | F | UF | F | UF | F | UF | F | UF |
| L | Posterior | .025 | .028 | .024 | .016 | .028 | .021 | .061 | .035 | .019 | .018 |
| R | | .027 | .028 | .024 | .017 | .030 | .028 | .053 | .024 | .020 | .019 |
| L | Central | .013 | .018 | .032 | .013 | .015 | .017 | .110 | .036 | .011 | .010 |
| R | | .016 | .010 | .027 | .010 | .011 | .012 | .074 | .025 | .011 | .006 |
| L | Frontal | .008 | .008 | .016 | .007 | .016 | .015 | .035 | .024 | .009 | .007 |
| R | | .008 | .008 | .012 | .009 | .011 | .019 | .031 | .013 | .007 | .009 |

**Supplementary Table 3.** Group differences in intersubject correlation values in the frequency bands, split by region. Significance of difference between the groups was assessed with a group assignment permutation test and a Bonferroni-Holm correction within a frequency band. Significant differences are denoted *** = $p < .001$, ** $p < .01$, * = $p < .05$.

| | | Familiar – Unfamiliar ISCs | | | | |
|---|---|---|---|---|---|---|
| Hemisphere | Region | Alpha | Beta | Delta | Gamma | Theta |
| L | Posterior | -.003 | .008*** | .006 | .025*** | .001 |
| R | | -.001 | .007*** | .002 | .029*** | .001 |
| L | Central | -.005 | .020*** | -.002 | .072*** | .001 |

| | | | | | | |
|---|---|---|---|---|---|---|
| R | | .006* | .020*** | -.001 | .049*** | .005 |
| L | Frontal | .000 | .009 | .002 | .012 | .002 |
| R | | .001 | .003 | -.007 | .018** | -.001 |

**Supplementary Table 4.** The EEG-fMRI correlations for each Schaefer 100 parcel for each frequency band across all participants. Values are Fisher's z transformed correlation values. Correlations were tested against 0 with a sign-flip permutation test and false discovery rate corrected within a frequency band. Significant differences are denoted *** = $p < .001$, ** $p < .01$, * = $p < .05$.

| Region | Alpha | Beta | Delta | Gamma | Theta |
|---|---|---|---|---|---|
| LH_Cont_Cing_1 | 0.013*** | 0.01*** | 0.016*** | 0.005*** | 0.008*** |
| LH_Cont_PFCl_1 | 0.009*** | 0.005*** | 0.008*** | 0.000 | 0.007*** |
| LH_Cont_Par_1 | -0.001 | 0.001 | 0.004** | 0.007*** | -0.002 |
| LH_Cont_pCun_1 | -0.01*** | 0.001 | 0.013*** | 0.01*** | 0.003* |
| LH_Default_PFC_1 | 0.005*** | 0.006*** | 0.003* | 0.002 | 0.001 |
| LH_Default_PFC_2 | 0.006*** | 0.004** | 0.003* | 0.000 | 0.003* |
| LH_Default_PFC_3 | 0.004** | 0.003* | 0.004*** | 0.000 | 0.000 |
| LH_Default_PFC_4 | 0.005*** | 0.003 | 0.007*** | -0.001 | 0.002 |
| LH_Default_PFC_5 | 0.004** | -0.001 | -0.001 | -0.004* | -0.002 |
| LH_Default_PFC_6 | 0.009*** | 0.005*** | 0.006*** | 0.000 | 0.005*** |
| LH_Default_PFC_7 | 0.007*** | 0.003 | 0.005*** | -0.001 | 0.002 |
| LH_Default_Par_1 | -0.014*** | 0.003* | -0.013*** | 0.01*** | -0.005*** |
| LH_Default_Par_2 | 0.008*** | 0.007*** | 0.007*** | 0.003 | 0.002* |
| LH_Default_Temp_1 | 0.003** | 0.004** | 0.001 | -0.002 | 0.004** |
| LH_Default_Temp_2 | 0.01*** | 0.012*** | 0.008*** | 0.003* | 0.013*** |
| LH_Default_pCunPCC_1 | -0.001 | 0.007*** | 0.018*** | 0.008*** | 0.008*** |
| LH_Default_pCunPCC_2 | 0.004** | 0.008*** | 0.01*** | 0.007*** | 0.006*** |
| LH_DorsAttn_FEF_1 | -0.012*** | -0.007*** | -0.012*** | 0.003 | -0.014*** |
| LH_DorsAttn_Post_1 | -0.034*** | -0.002 | -0.018*** | 0.026*** | -0.019*** |
| LH_DorsAttn_Post_2 | -0.019*** | -0.016*** | -0.021*** | -0.003 | -0.021*** |
| LH_DorsAttn_Post_3 | -0.023*** | -0.011*** | -0.01*** | 0.011*** | -0.017*** |
| LH_DorsAttn_Post_4 | -0.019*** | -0.009*** | -0.019*** | 0.007*** | -0.022*** |
| LH_DorsAttn_Post_5 | -0.024*** | -0.002 | -0.005*** | 0.02*** | -0.011*** |
| LH_DorsAttn_Post_6 | -0.034*** | -0.02*** | -0.026*** | 0.006*** | -0.032*** |
| LH_DorsAttn_PrCv_1 | -0.003** | -0.001 | -0.005*** | 0.002 | -0.003* |
| LH_Limbic_OFC_1 | -0.001 | 0.003* | 0.001 | 0.004* | -0.001 |
| LH_Limbic_TempPole_1 | -0.005*** | 0.001 | -0.005*** | 0.004** | -0.004*** |

| | | | | | |
|---|---|---|---|---|---|
| LH_Limbic_TempPole_2 | -0.002 | -0.002 | 0.000 | -0.001 | -0.002 |
| LH_SalVentAttn_FrOperIns_1 | 0.000 | -0.001 | 0.001 | -0.002 | 0.000 |
| LH_SalVentAttn_FrOperIns_2 | 0.009*** | 0.004** | 0.01*** | -0.003* | 0.005*** |
| LH_SalVentAttn_Med_1 | 0.013*** | 0.007*** | 0.013*** | -0.001 | 0.006*** |
| LH_SalVentAttn_Med_2 | -0.015*** | -0.006*** | -0.007*** | 0.006*** | -0.013*** |
| LH_SalVentAttn_Med_3 | 0.003* | 0.002 | 0.004** | -0.001 | -0.001 |
| LH_SalVentAttn_PFCl_1 | 0.01*** | 0.006*** | 0.008*** | 0.000 | 0.004*** |
| LH_SalVentAttn_ParOper_1 | -0.021*** | -0.012*** | -0.019*** | 0.002 | -0.019*** |
| LH_SomMot_1 | -0.016*** | -0.003* | -0.012*** | 0.007*** | -0.004** |
| LH_SomMot_2 | 0.004*** | 0.001 | 0.001 | -0.005** | 0.000 |
| LH_SomMot_3 | -0.009*** | -0.004** | -0.007*** | 0.002 | -0.007*** |
| LH_SomMot_4 | 0.004*** | 0.002 | 0.001 | -0.003 | -0.001 |
| LH_SomMot_5 | 0.004** | 0.003 | -0.001 | 0.000 | -0.004** |
| LH_SomMot_6 | -0.005*** | -0.002 | -0.002 | 0.000 | -0.008*** |
| LH_Vis_1 | -0.003* | 0.005*** | 0.012*** | 0.01*** | 0.004** |
| LH_Vis_2 | -0.036*** | 0.01*** | -0.001 | 0.045*** | -0.006*** |
| LH_Vis_3 | -0.033*** | -0.006*** | -0.001 | 0.02*** | -0.009*** |
| LH_Vis_4 | -0.013*** | 0.027*** | 0.014*** | 0.049*** | 0.005*** |
| LH_Vis_5 | -0.037*** | 0.011*** | -0.005** | 0.048*** | -0.008*** |
| LH_Vis_6 | -0.023*** | 0.000 | 0.009*** | 0.014*** | -0.001 |
| LH_Vis_7 | -0.066*** | -0.023*** | -0.046*** | 0.028*** | -0.043*** |
| LH_Vis_8 | -0.053*** | -0.012*** | -0.021*** | 0.034*** | -0.027*** |
| LH_Vis_9 | -0.031*** | 0.001 | 0.008*** | 0.025*** | -0.002 |
| RH_Cont_Cing_1 | 0.012*** | 0.009*** | 0.017*** | 0.005*** | 0.008*** |
| RH_Cont_PFCl_1 | 0.005*** | 0.002 | 0.006*** | 0.000 | 0.003** |
| RH_Cont_PFCl_2 | 0.01*** | 0.007*** | 0.012*** | 0.001 | 0.01*** |
| RH_Cont_PFCl_3 | 0.009*** | 0.006*** | 0.01*** | 0.000 | 0.007*** |
| RH_Cont_PFCl_4 | 0.003* | 0.001 | 0.003* | -0.001 | 0.004** |
| RH_Cont_PFCmp_1 | 0.015*** | 0.007*** | 0.013*** | -0.004* | 0.008*** |
| RH_Cont_Par_1 | -0.004** | -0.002 | -0.004** | 0.003* | -0.007*** |
| RH_Cont_Par_2 | 0.012*** | 0.008*** | 0.017*** | 0.004* | 0.01*** |
| RH_Cont_pCun_1 | -0.015*** | 0.000 | 0.006*** | 0.013*** | -0.001 |
| RH_Default_PFCdPFCm_1 | 0.001 | 0.000 | 0.001 | -0.003 | -0.001 |
| RH_Default_PFCdPFCm_2 | -0.004** | -0.005** | -0.006*** | -0.005* | -0.005*** |
| RH_Default_PFCdPFCm_3 | 0.002 | -0.001 | 0.004*** | -0.002 | 0.002 |
| RH_Default_PFCv_1 | 0.002* | 0.006*** | 0.004*** | 0.005** | 0.001 |
| RH_Default_PFCv_2 | 0.000 | 0.001 | -0.003* | -0.001 | -0.001 |
| RH_Default_Par_1 | -0.007*** | 0.001 | -0.006*** | 0.006*** | -0.006*** |

| | | | | | |
|---|---|---|---|---|---|
| RH_Default_Temp_1 | 0.001 | 0.006*** | 0.005*** | 0.008*** | 0.003* |
| RH_Default_Temp_2 | -0.001 | 0.014*** | 0.005** | 0.011*** | 0.009*** |
| RH_Default_Temp_3 | -0.005*** | 0.011*** | -0.002 | 0.011*** | 0.006*** |
| RH_Default_pCunPCC_1 | -0.004*** | 0.006*** | 0.018*** | 0.009*** | 0.007*** |
| RH_Default_pCunPCC_2 | -0.001 | 0.007*** | 0.008*** | 0.009*** | 0.005*** |
| RH_DorsAttn_FEF_1 | -0.016*** | -0.01*** | -0.012*** | 0.001 | -0.014*** |
| RH_DorsAttn_Post_1 | -0.042*** | -0.012*** | -0.024*** | 0.02*** | -0.026*** |
| RH_DorsAttn_Post_2 | -0.009*** | -0.006*** | -0.013*** | 0.002 | -0.013*** |
| RH_DorsAttn_Post_3 | -0.011*** | -0.006*** | -0.005*** | 0.007*** | -0.009*** |
| RH_DorsAttn_Post_4 | -0.024*** | -0.01*** | -0.005*** | 0.012*** | -0.014*** |
| RH_DorsAttn_Post_5 | -0.038*** | -0.016*** | -0.019*** | 0.012*** | -0.027*** |
| RH_DorsAttn_PrCv_1 | -0.007*** | 0.002 | -0.005*** | 0.005** | -0.003* |
| RH_Limbic_OFC_1 | -0.002 | 0.001 | 0.001 | 0.002 | -0.001 |
| RH_Limbic_TempPole_1 | -0.007*** | 0.003* | -0.003* | 0.009*** | -0.004** |
| RH_SalVentAttn_FrOperIns_1 | 0.005*** | 0.003* | 0.007*** | -0.003* | 0.003* |
| RH_SalVentAttn_Med_1 | -0.014*** | -0.006*** | -0.005*** | 0.005** | -0.012*** |
| RH_SalVentAttn_Med_2 | 0.001 | 0.003 | 0.004*** | 0.002 | -0.002 |
| RH_SalVentAttn_TempOccPar_1 | -0.026*** | 0.000 | -0.018*** | 0.015*** | -0.012*** |
| RH_SalVentAttn_TempOccPar_2 | -0.025*** | -0.014*** | -0.024*** | 0.003 | -0.023*** |
| RH_SomMot_1 | -0.018*** | -0.003* | -0.013*** | 0.008*** | -0.007*** |
| RH_SomMot_2 | 0.006*** | 0.004** | 0.001 | -0.002 | 0.000 |
| RH_SomMot_3 | -0.003* | -0.001 | -0.004** | 0.000 | -0.002* |
| RH_SomMot_4 | 0.006*** | 0.004** | 0.001 | -0.001 | 0.000 |
| RH_SomMot_5 | 0.002* | 0.001 | -0.001 | -0.003 | -0.002 |
| RH_SomMot_6 | 0.005*** | 0.003 | -0.001 | -0.001 | -0.001 |
| RH_SomMot_7 | -0.023*** | -0.014*** | -0.019*** | 0.004* | -0.024*** |
| RH_SomMot_8 | -0.004** | -0.001 | 0.000 | 0.000 | -0.006*** |
| RH_Vis_1 | -0.012*** | 0.008*** | 0.01*** | 0.02*** | 0.003* |
| RH_Vis_2 | -0.042*** | -0.002 | -0.007*** | 0.033*** | -0.013*** |
| RH_Vis_3 | -0.043*** | 0.001 | -0.018*** | 0.035*** | -0.022*** |
| RH_Vis_4 | -0.021*** | 0.022*** | 0.012*** | 0.048*** | 0.000 |
| RH_Vis_5 | -0.045*** | -0.005*** | -0.016*** | 0.031*** | -0.019*** |
| RH_Vis_6 | -0.02*** | -0.004** | 0.008*** | 0.009*** | -0.002 |
| RH_Vis_7 | -0.054*** | -0.015*** | -0.022*** | 0.029*** | -0.03*** |
| RH_Vis_8 | -0.053*** | -0.009*** | -0.011*** | 0.032*** | -0.02*** |

**Supplementary Table 5.** The EEG-fMRI correlations within each Yeo (2011) network for each frequency band across all participants. Values are Fisher's z transformed correlation values. Correlations were tested against 0 with a sign-flip permutation test and false discovery rate corrected within a frequency band. Significant differences are denoted *** = $p < .001$, ** $p < .01$, * = $p < .05$.

| Network | Alpha | Beta | Delta | Gamma | Theta |
|---|---|---|---|---|---|
| Visual | -.034*** | .001 | -.005*** | .030*** | -.011*** |
| Dorsal Attention | -.021*** | -.009*** | -.013*** | .009*** | -.017*** |
| Ventral Attention | -.005*** | -.001 | -.002* | .002 | -.005*** |
| Default Mode | .001 | .004*** | .004*** | .003** | .002** |
| Somato-motor | -.003** | -.001 | -.004*** | .001 | -.005*** |
| Fronto-parietal | .004*** | .004*** | .009*** | .003** | .005*** |
| Limbic | -.003** | .001 | -.001 | .004** | -.002* |

**Supplementary Table 6.** The EEG-fMRI correlations for each Schaefer 100 parcel for each frequency band in familiar participants. Values are Fisher's z transformed correlation values. Correlations were tested against 0 with a sign-flip permutation test and false discovery rate corrected within a frequency band. Significant differences are denoted *** = $p < .001$, ** $p < .01$, * = $p < .05$.

| Region | Alpha | Beta | Delta | Gamma | Theta |
|---|---|---|---|---|---|
| LH_Cont_Cing_1 | 0.012*** | 0.007* | 0.01*** | 0.002 | 0.002 |
| LH_Cont_PFCl_1 | 0.007* | -0.003 | 0.003 | -0.006 | 0.006* |
| LH_Cont_Par_1 | -0.003 | -0.008* | -0.002 | -0.001 | -0.005* |
| LH_Cont_pCun_1 | -0.01*** | -0.001 | 0.003 | 0.009** | -0.001 |
| LH_Default_PFC_1 | 0.005 | 0.006* | 0.003 | 0.004 | 0.003 |
| LH_Default_PFC_2 | 0.007** | 0.006 | 0.004 | 0.001 | 0.005 |
| LH_Default_PFC_3 | 0.003 | 0.003 | -0.001 | 0.002 | -0.004 |
| LH_Default_PFC_4 | 0.002 | 0.001 | 0.003 | 0.003 | -0.005 |
| LH_Default_PFC_5 | 0.005 | 0.000 | -0.002 | -0.006 | -0.003 |
| LH_Default_PFC_6 | 0.008** | 0.004 | 0.003 | 0.000 | 0.006* |
| LH_Default_PFC_7 | 0.004 | -0.001 | 0.000 | -0.003 | -0.002 |
| LH_Default_Par_1 | -0.009** | 0.018*** | -0.005 | 0.022*** | 0.006* |
| LH_Default_Par_2 | 0.009** | 0.008** | 0.004 | 0.003 | 0.003 |
| LH_Default_Temp_1 | 0.008** | 0.009** | 0.004 | -0.001 | 0.012*** |
| LH_Default_Temp_2 | 0.015*** | 0.022*** | 0.012*** | 0.009** | 0.02*** |
| LH_Default_pCunPCC_1 | -0.001 | 0.009** | 0.013*** | 0.011*** | 0.009*** |

| | | | | | |
|---|---|---|---|---|---|
| LH_Default_pCunPCC_2 | 0.005 | 0.014*** | 0.006* | 0.013*** | 0.009*** |
| LH_DorsAttn_FEF_1 | -0.014*** | -0.016*** | -0.017*** | -0.005 | -0.014*** |
| LH_DorsAttn_Post_1 | -0.035*** | 0.003 | -0.017*** | 0.031*** | -0.014*** |
| LH_DorsAttn_Post_2 | -0.025*** | -0.026*** | -0.025*** | -0.008* | -0.024*** |
| LH_DorsAttn_Post_3 | -0.022*** | -0.017*** | -0.015*** | 0.007 | -0.014*** |
| LH_DorsAttn_Post_4 | -0.024*** | -0.015*** | -0.022*** | 0.007* | -0.025*** |
| LH_DorsAttn_Post_5 | -0.026*** | 0.000 | -0.012*** | 0.021*** | -0.01*** |
| LH_DorsAttn_Post_6 | -0.038*** | -0.026*** | -0.033*** | 0.002 | -0.034*** |
| LH_DorsAttn_PrCv_1 | -0.006 | -0.006 | -0.007** | 0.000 | -0.001 |
| LH_Limbic_OFC_1 | -0.002 | 0.011*** | 0.000 | 0.018*** | -0.002 |
| LH_Limbic_TempPole_1 | -0.003 | 0.004 | -0.002 | 0.006 | 0.003 |
| LH_Limbic_TempPole_2 | -0.005 | -0.008** | -0.004 | -0.006 | -0.004 |
| LH_SalVentAttn_FrOperIns_1 | 0.001 | -0.009** | -0.003 | -0.011*** | 0.002 |
| LH_SalVentAttn_FrOperIns_2 | 0.006* | -0.006* | 0.006* | -0.013*** | 0.002 |
| LH_SalVentAttn_Med_1 | 0.01*** | -0.003 | 0.006* | -0.009* | 0.001 |
| LH_SalVentAttn_Med_2 | -0.017*** | -0.011** | -0.015*** | 0.001 | -0.015*** |
| LH_SalVentAttn_Med_3 | 0.001 | -0.005 | 0.000 | -0.009* | -0.004 |
| LH_SalVentAttn_PFCl_1 | 0.007* | 0.000 | 0.004 | -0.004 | 0.000 |
| LH_SalVentAttn_ParOper_1 | -0.021*** | -0.012*** | -0.019*** | 0.001 | -0.017*** |
| LH_SomMot_1 | -0.01*** | 0.005 | -0.012*** | 0.01*** | 0.000 |
| LH_SomMot_2 | 0.002 | -0.006 | -0.006* | -0.009** | -0.004 |
| LH_SomMot_3 | -0.008*** | -0.008** | -0.012*** | -0.003 | -0.007** |
| LH_SomMot_4 | 0.000 | -0.006 | -0.005* | -0.007 | -0.002 |
| LH_SomMot_5 | 0.001 | -0.005 | -0.003 | -0.006 | -0.006* |
| LH_SomMot_6 | -0.008** | -0.009* | -0.007** | -0.007 | -0.012*** |
| LH_Vis_1 | -0.003 | 0.001 | 0.008** | 0.006* | 0.005* |
| LH_Vis_2 | -0.033*** | 0.02*** | -0.002 | 0.051*** | -0.001 |
| LH_Vis_3 | -0.03*** | -0.002 | -0.005 | 0.022*** | -0.008** |
| LH_Vis_4 | -0.011*** | 0.032*** | 0.014*** | 0.05*** | 0.008** |
| LH_Vis_5 | -0.035*** | 0.021*** | -0.006 | 0.051*** | -0.006* |
| LH_Vis_6 | -0.018 | 0.003 | 0.005 | 0.014*** | -0.001 |
| LH_Vis_7 | -0.066*** | -0.016*** | -0.042*** | 0.033*** | -0.038*** |
| LH_Vis_8 | -0.051*** | -0.007* | -0.021*** | 0.038*** | -0.022*** |
| LH_Vis_9 | -0.028 | 0.004 | -0.001 | 0.024*** | -0.003 |
| RH_Cont_Cing_1 | 0.011*** | 0.004 | 0.011*** | 0.001 | 0.003 |
| RH_Cont_PFCl_1 | 0.002 | 0.001 | 0.002 | 0.005 | 0.000 |
| RH_Cont_PFCl_2 | 0.004 | 0.002 | 0.007** | 0.003 | 0.007** |
| RH_Cont_PFCl_3 | 0.005 | 0.002 | 0.005 | -0.001 | 0.003 |

| | | | | | |
|---|---|---|---|---|---|
| RH_Cont_PFCl_4 | 0.000 | 0.003 | 0.000 | 0.004 | 0.004 |
| RH_Cont_PFCmp_1 | 0.011*** | 0.000 | 0.008*** | -0.007* | 0.004 |
| RH_Cont_Par_1 | -0.003 | -0.006 | -0.006* | -0.001 | -0.009*** |
| RH_Cont_Par_2 | 0.01*** | 0.005 | 0.012*** | 0.001 | 0.008** |
| RH_Cont_pCun_1 | -0.018 | 0.002 | -0.001 | 0.016*** | 0.000 |
| RH_Default_PFCdPFCm_1 | 0.000 | 0.001 | -0.002 | 0.001 | -0.002 |
| RH_Default_PFCdPFCm_2 | -0.002 | -0.001 | -0.006* | -0.002 | -0.002 |
| RH_Default_PFCdPFCm_3 | -0.002 | -0.002 | -0.001 | -0.001 | 0.000 |
| RH_Default_PFCv_1 | 0.003 | 0.01*** | 0.002 | 0.014*** | 0.001 |
| RH_Default_PFCv_2 | 0.002 | 0.011*** | 0.003 | 0.013*** | 0.006* |
| RH_Default_Par_1 | -0.005 | 0.008** | -0.005* | 0.013*** | -0.003 |
| RH_Default_Temp_1 | 0.002 | 0.018*** | 0.004 | 0.024*** | 0.005 |
| RH_Default_Temp_2 | 0.004 | 0.027*** | 0.01*** | 0.024*** | 0.019*** |
| RH_Default_Temp_3 | 0.003 | 0.028*** | 0.004 | 0.025*** | 0.014*** |
| RH_Default_pCunPCC_1 | -0.003 | 0.011*** | 0.013*** | 0.015*** | 0.01*** |
| RH_Default_pCunPCC_2 | -0.001 | 0.014*** | 0.005 | 0.017*** | 0.008** |
| RH_DorsAttn_FEF_1 | -0.019*** | -0.018*** | -0.016*** | -0.005 | -0.014*** |
| RH_DorsAttn_Post_1 | -0.04*** | -0.004 | -0.023*** | 0.027*** | -0.018*** |
| RH_DorsAttn_Post_2 | -0.011*** | -0.011*** | -0.015*** | 0.002 | -0.015*** |
| RH_DorsAttn_Post_3 | -0.014*** | -0.015*** | -0.01*** | 0.000 | -0.012*** |
| RH_DorsAttn_Post_4 | -0.025*** | -0.015*** | -0.01*** | 0.009* | -0.012*** |
| RH_DorsAttn_Post_5 | -0.04*** | -0.019*** | -0.025*** | 0.009* | -0.026*** |
| RH_DorsAttn_PrCv_1 | -0.008** | 0.005 | -0.005 | 0.014*** | 0.002 |
| RH_Limbic_OFC_1 | -0.003 | 0.011** | 0.002 | 0.017*** | -0.002 |
| RH_Limbic_TempPole_1 | -0.005 | 0.016*** | -0.001 | 0.025*** | 0.002 |
| RH_SalVentAttn_FrOperIns_1 | 0.002 | -0.005 | 0.000 | -0.006* | -0.002 |
| RH_SalVentAttn_Med_1 | -0.016*** | -0.013*** | -0.013*** | -0.001 | -0.017*** |
| RH_SalVentAttn_Med_2 | -0.002 | -0.005 | 0.000 | -0.007 | -0.005* |
| RH_SalVentAttn_TempOccPar_1 | -0.022*** | 0.011*** | -0.015*** | 0.021*** | -0.004 |
| RH_SalVentAttn_TempOccPar_2 | -0.028*** | -0.016*** | -0.026*** | 0.005 | -0.026*** |
| RH_SomMot_1 | -0.014*** | 0.001 | -0.018*** | 0.012*** | -0.009*** |
| RH_SomMot_2 | 0.003 | -0.004 | -0.008** | -0.006 | -0.006** |
| RH_SomMot_3 | -0.003 | -0.003 | -0.01*** | 0.000 | -0.006* |
| RH_SomMot_4 | 0.003 | 0.002 | -0.005 | 0.001 | -0.003 |
| RH_SomMot_5 | 0.002 | -0.002 | -0.005 | -0.004 | -0.003 |
| RH_SomMot_6 | 0.000 | -0.005 | -0.005* | -0.007 | -0.004 |
| RH_SomMot_7 | -0.024*** | -0.015*** | -0.022*** | 0.003 | -0.026*** |
| RH_SomMot_8 | -0.008** | -0.008* | -0.006* | -0.007 | -0.01*** |

| | | | | | |
|---|---|---|---|---|---|
| RH_Vis_1 | -0.013*** | 0.008** | 0.003 | 0.022*** | 0.002 |
| RH_Vis_2 | -0.039*** | 0.01** | -0.008** | 0.044*** | -0.009** |
| RH_Vis_3 | -0.043*** | 0.011*** | -0.019*** | 0.044*** | -0.02*** |
| RH_Vis_4 | -0.015 | 0.037*** | 0.015*** | 0.058*** | 0.005 |
| RH_Vis_5 | -0.039*** | 0.006* | -0.019*** | 0.037*** | -0.018*** |
| RH_Vis_6 | -0.014 | 0.002 | 0.002 | 0.013*** | -0.002 |
| RH_Vis_7 | -0.051*** | -0.009** | -0.021*** | 0.036*** | -0.026*** |
| RH_Vis_8 | -0.049*** | 0.001 | -0.014*** | 0.038*** | -0.018*** |

**Supplementary Table 7.** The EEG-fMRI correlations for each Schaefer 100 parcel for each frequency band in unfamiliar participants. Values are Fisher's z transformed correlation values. Correlations were tested against 0 with a sign-flip permutation test and false discovery rate corrected within a frequency band. Significant differences are denoted *** = $p < .001$, ** $p < .01$, * = $p < .05$.

| Region | Alpha | Beta | Delta | Gamma | Theta |
|---|---|---|---|---|---|
| LH_Cont_Cing_1 | 0.015*** | 0.014*** | 0.024*** | 0.01** | 0.017*** |
| LH_Cont_PFCl_1 | 0.011*** | 0.011*** | 0.014*** | 0.005 | 0.01** |
| LH_Cont_Par_1 | 0.001 | 0.008* | 0.013*** | 0.013*** | 0.004 |
| LH_Cont_pCun_1 | -0.01*** | 0.002 | 0.025*** | 0.013*** | 0.01*** |
| LH_Default_PFC_1 | 0.003 | 0.008** | 0.003 | 0.002 | 0.000 |
| LH_Default_PFC_2 | 0.004 | 0.005 | 0.003 | -0.001 | 0.002 |
| LH_Default_PFC_3 | 0.005 | 0.008** | 0.011*** | 0.002 | 0.003 |
| LH_Default_PFC_4 | 0.007** | 0.005 | 0.012*** | -0.004 | 0.007** |
| LH_Default_PFC_5 | 0.002 | 0.001 | 0.001 | 0.000 | -0.002 |
| LH_Default_PFC_6 | 0.009*** | 0.008* | 0.011*** | 0.002 | 0.008** |
| LH_Default_PFC_7 | 0.01*** | 0.008* | 0.012*** | 0.003 | 0.006* |
| LH_Default_Par_1 | -0.017*** | -0.007* | -0.019*** | 0.004 | -0.013*** |
| LH_Default_Par_2 | 0.006* | 0.007* | 0.013*** | 0.004 | 0.005 |
| LH_Default_Temp_1 | -0.003 | 0.001 | 0.000 | 0.000 | -0.001 |
| LH_Default_Temp_2 | 0.005 | 0.008** | 0.006* | 0.002 | 0.009*** |
| LH_Default_pCunPCC_1 | -0.003 | 0.007** | 0.026*** | 0.01*** | 0.008** |
| LH_Default_pCunPCC_2 | 0.002 | 0.006* | 0.016*** | 0.007* | 0.007* |
| LH_DorsAttn_FEF_1 | -0.01*** | -0.002 | -0.006* | 0.011* | -0.012*** |
| LH_DorsAttn_Post_1 | -0.032*** | -0.005 | -0.016*** | 0.026*** | -0.02*** |
| LH_DorsAttn_Post_2 | -0.014*** | -0.01*** | -0.014*** | 0.002 | -0.016*** |
| LH_DorsAttn_Post_3 | -0.022*** | -0.006 | -0.002 | 0.016*** | -0.015*** |

| | | | | | |
|---|---|---|---|---|---|
| LH_DorsAttn_Post_4 | -0.013*** | -0.006* | -0.013*** | 0.008* | -0.015*** |
| LH_DorsAttn_Post_5 | -0.022*** | -0.004 | 0.004 | 0.021*** | -0.01*** |
| LH_DorsAttn_Post_6 | -0.028*** | -0.015*** | -0.018*** | 0.011** | -0.027*** |
| LH_DorsAttn_PrCv_1 | -0.001 | 0.002 | -0.002 | 0.004 | -0.003 |
| LH_Limbic_OFC_1 | 0.000 | -0.001 | 0.005 | -0.007* | 0.002 |
| LH_Limbic_TempPole_1 | -0.008** | -0.001 | -0.003 | 0.003 | -0.007** |
| LH_Limbic_TempPole_2 | 0.002 | 0.004 | 0.007* | 0.006 | 0.005 |
| LH_SalVentAttn_FrOperIns_1 | -0.001 | 0.005 | 0.006* | 0.007* | -0.001 |
| LH_SalVentAttn_FrOperIns_2 | 0.01*** | 0.01*** | 0.016*** | 0.005 | 0.008** |
| LH_SalVentAttn_Med_1 | 0.015*** | 0.014*** | 0.02*** | 0.007 | 0.012*** |
| LH_SalVentAttn_Med_2 | -0.013*** | -0.002 | 0.004 | 0.012*** | -0.007* |
| LH_SalVentAttn_Med_3 | 0.003 | 0.006 | 0.008** | 0.008 | 0.001 |
| LH_SalVentAttn_PFCl_1 | 0.011*** | 0.009** | 0.013*** | 0.006 | 0.01** |
| LH_SalVentAttn_ParOper_1 | -0.022*** | -0.015*** | -0.018*** | 0.003 | -0.019*** |
| LH_SomMot_1 | -0.022*** | -0.008** | -0.01*** | 0.006** | -0.005* |
| LH_SomMot_2 | 0.006** | 0.007** | 0.012*** | 0.000 | 0.005 |
| LH_SomMot_3 | -0.008** | -0.001 | 0.001 | 0.007* | -0.003 |
| LH_SomMot_4 | 0.011*** | 0.009*** | 0.01*** | 0.002 | 0.003 |
| LH_SomMot_5 | 0.008** | 0.009** | 0.007* | 0.008* | 0.003 |
| LH_SomMot_6 | 0.000 | 0.003 | 0.009** | 0.007 | 0.000 |
| LH_Vis_1 | -0.003 | 0.009*** | 0.018*** | 0.016*** | 0.003 |
| LH_Vis_2 | -0.039*** | 0.001 | 0.000 | 0.044*** | -0.01** |
| LH_Vis_3 | -0.036*** | -0.011*** | 0.005 | 0.019*** | -0.011*** |
| LH_Vis_4 | -0.013*** | 0.024*** | 0.012*** | 0.051*** | 0.004 |
| LH_Vis_5 | -0.038*** | 0.002 | -0.004 | 0.047*** | -0.009** |
| LH_Vis_6 | -0.028*** | -0.003 | 0.016*** | 0.016*** | -0.002 |
| LH_Vis_7 | -0.066*** | -0.029*** | -0.049*** | 0.025*** | -0.048*** |
| LH_Vis_8 | -0.055*** | -0.015*** | -0.02*** | 0.034*** | -0.03*** |
| LH_Vis_9 | -0.033*** | -0.002 | 0.019*** | 0.03*** | 0.002 |
| RH_Cont_Cing_1 | 0.014*** | 0.015*** | 0.024*** | 0.011*** | 0.016*** |
| RH_Cont_PFCl_1 | 0.005 | 0.003 | 0.012*** | -0.005 | 0.008** |
| RH_Cont_PFCl_2 | 0.015*** | 0.013*** | 0.015*** | 0.000 | 0.012*** |
| RH_Cont_PFCl_3 | 0.011*** | 0.007* | 0.016*** | 0.001 | 0.013*** |
| RH_Cont_PFCl_4 | 0.004 | 0.000 | 0.005 | -0.006 | 0.003 |
| RH_Cont_PFCmp_1 | 0.017*** | 0.012*** | 0.019*** | 0.000 | 0.013*** |
| RH_Cont_Par_1 | -0.006* | -0.003 | -0.002 | 0.003 | -0.005 |
| RH_Cont_Par_2 | 0.01*** | 0.01*** | 0.023*** | 0.007* | 0.012*** |
| RH_Cont_pCun_1 | -0.013*** | -0.002 | 0.017*** | 0.013*** | 0.004 |

| | | | | | |
|---|---|---|---|---|---|
| RH_Default_PFCdPFCm_1 | 0.001 | 0.001 | 0.005 | -0.003 | 0.000 |
| RH_Default_PFCdPFCm_2 | -0.006* | -0.009** | -0.008** | -0.006 | -0.009*** |
| RH_Default_PFCdPFCm_3 | 0.004 | 0.001 | 0.01*** | -0.002 | 0.004 |
| RH_Default_PFCv_1 | 0.003 | 0.006 | 0.005 | -0.003 | 0.001 |
| RH_Default_PFCv_2 | -0.001 | -0.006 | -0.009** | -0.014*** | -0.008** |
| RH_Default_Par_1 | -0.01*** | -0.008** | -0.008** | -0.001 | -0.009*** |
| RH_Default_Temp_1 | 0.001 | -0.001 | 0.008** | -0.004 | 0.004 |
| RH_Default_Temp_2 | -0.006* | 0.002 | -0.001 | 0.000 | -0.001 |
| RH_Default_Temp_3 | -0.011*** | -0.003 | -0.009** | 0.000 | -0.002 |
| RH_Default_pCunPCC_1 | -0.007** | 0.002 | 0.025*** | 0.007** | 0.004 |
| RH_Default_pCunPCC_2 | 0.000 | 0.004 | 0.013*** | 0.005* | 0.004 |
| RH_DorsAttn_FEF_1 | -0.015*** | -0.009** | -0.007** | 0.005 | -0.014*** |
| RH_DorsAttn_Post_1 | -0.044*** | -0.02*** | -0.026*** | 0.014*** | -0.033*** |
| RH_DorsAttn_Post_2 | -0.008** | -0.005 | -0.01*** | 0.001 | -0.011*** |
| RH_DorsAttn_Post_3 | -0.01*** | -0.002 | 0.001 | 0.011** | -0.006* |
| RH_DorsAttn_Post_4 | -0.024*** | -0.007* | 0.002 | 0.015*** | -0.013*** |
| RH_DorsAttn_Post_5 | -0.036*** | -0.017*** | -0.01*** | 0.015*** | -0.025*** |
| RH_DorsAttn_PrCv_1 | -0.006* | -0.003 | -0.007* | -0.004 | -0.007** |
| RH_Limbic_OFC_1 | -0.001 | -0.005 | 0.003 | -0.013*** | 0.001 |
| RH_Limbic_TempPole_1 | -0.006* | -0.005 | -0.004 | -0.005 | -0.007** |
| RH_SalVentAttn_FrOperIns_1 | 0.008** | 0.009** | 0.014*** | 0.000 | 0.007* |
| RH_SalVentAttn_Med_1 | -0.012*** | -0.002 | 0.006* | 0.01** | -0.006* |
| RH_SalVentAttn_Med_2 | 0.004 | 0.008* | 0.01*** | 0.01* | 0.001 |
| RH_SalVentAttn_TempOccPar_1 | -0.028*** | -0.009** | -0.022*** | 0.01*** | -0.019*** |
| RH_SalVentAttn_TempOccPar_2 | -0.021*** | -0.013*** | -0.019*** | 0.002 | -0.019*** |
| RH_SomMot_1 | -0.02*** | -0.005 | -0.006* | 0.007*** | -0.005 |
| RH_SomMot_2 | 0.009*** | 0.01*** | 0.012*** | 0.001 | 0.005 |
| RH_SomMot_3 | -0.001 | 0.003 | 0.006* | 0.002 | 0.003 |
| RH_SomMot_4 | 0.01*** | 0.008** | 0.011*** | -0.001 | 0.003 |
| RH_SomMot_5 | 0.005* | 0.005 | 0.006* | 0.001 | 0.000 |
| RH_SomMot_6 | 0.01*** | 0.01*** | 0.005 | 0.007 | 0.004 |
| RH_SomMot_7 | -0.022*** | -0.015*** | -0.015*** | 0.005 | -0.02*** |
| RH_SomMot_8 | 0.001 | 0.006 | 0.009** | 0.01* | 0.001 |
| RH_Vis_1 | -0.011*** | 0.007** | 0.014*** | 0.018*** | 0.001 |
| RH_Vis_2 | -0.047*** | -0.013*** | -0.007* | 0.026*** | -0.017*** |
| RH_Vis_3 | -0.043*** | -0.007* | -0.019*** | 0.03*** | -0.023*** |
| RH_Vis_4 | -0.025*** | 0.01*** | 0.009** | 0.043*** | -0.004 |
| RH_Vis_5 | -0.051*** | -0.016*** | -0.012*** | 0.026*** | -0.02*** |

| | | | | | |
|---|---|---|---|---|---|
| RH_Vis_6 | -0.027*** | -0.01*** | 0.018*** | 0.007** | -0.001 |
| RH_Vis_7 | -0.056*** | -0.019*** | -0.024*** | 0.026*** | -0.035*** |
| RH_Vis_8 | -0.056*** | -0.017*** | -0.007* | 0.031*** | -0.02*** |

**Supplementary Table 8.** The EEG-fMRI correlations within each Yeo (2011) network for each frequency band in the familiar participants. Values are Fisher's z transformed correlation values. Correlations were tested against 0 with a sign-flip permutation test and false discovery rate corrected within a frequency band. Significant differences are denoted *** = *p* < .001, ** *p* < .01, * = *p* < .05.

| Network | Alpha | Beta | Delta | Gamma | Theta |
|---|---|---|---|---|---|
| Visual | -.032*** | .007** | -.006** | .034*** | -.009*** |
| Dorsal Attention | -.023*** | -.012** | -.017*** | .007* | -.015*** |
| Ventral Attention | -.007** | -.006* | -.006** | -.003 | -.007*** |
| Default Mode | .002 | .009** | .003 | .008** | .005** |
| Somato-motor | -.005 | -.004 | -.009*** | -.002 | -.007** |
| Fronto-parietal | .002 | .001 | .004 | .002 | .002 |
| Limbic | -.004 | .007** | -.001 | .012*** | -.000 |

**Supplementary Table 9.** The EEG-fMRI correlations within each Yeo (2011) network for each frequency band in the unfamiliar participants. Values are Fisher's z transformed correlation values. Correlations were tested against 0 with a sign-flip permutation test and false discovery rate corrected within a frequency band. Significant differences are denoted *** = *p* < .001, ** *p* < .01, * = *p* < .05.

| Network | Alpha | Beta | Delta | Gamma | Theta |
|---|---|---|---|---|---|
| Visual | -.037*** | -.005* | -.002 | .029*** | -.013*** |
| Dorsal Attention | -.019*** | -.007** | -.008*** | .010*** | -.015*** |
| Ventral Attention | -.004 | .002 | .003 | .006* | -.003 |
| Default Mode | .000 | .002 | .005* | .000 | .001 |
| Somato-motor | -.000 | .003 | .004 | .006* | -.000 |
| Fronto-parietal | .006** | .006** | .016*** | .005 | .009*** |
| Limbic | -.003 | -.002 | .001 | -.003 | -.001 |

**Supplementary Table 10.** The EEG-fMRI correlations for each Schaefer 100 parcel for each frequency band compared between familiar and unfamiliar participants. Values are Fisher's z transformed correlation difference values. Significance of difference between the groups was assessed with a group assignment permutation test and a false discovery rate correction within a frequency band. Significant differences are denoted *** = $p < .001$, ** $p < .01$, * = $p < .05$.

| | Familiar - Unfamiliar | | | | |
|---|---|---|---|---|---|
| **Region** | **Alpha** | **Beta** | **Delta** | **Gamma** | **Theta** |
| LH_Cont_Cing_1 | -0.002 | -0.007 | -0.014*** | -0.008 | -0.014*** |
| LH_Cont_PFCl_1 | -0.005 | -0.014** | -0.012** | -0.012* | -0.005 |
| LH_Cont_Par_1 | -0.004 | -0.016*** | -0.015*** | -0.014** | -0.009* |
| LH_Cont_pCun_1 | 0 | -0.002 | -0.022*** | -0.005 | -0.011** |
| LH_Default_PFC_1 | 0.002 | -0.002 | 0.000 | 0.003 | 0.003 |
| LH_Default_PFC_2 | 0.004 | 0.001 | 0.000 | 0.002 | 0.003 |
| LH_Default_PFC_3 | -0.002 | -0.005 | -0.011** | 0.000 | -0.007 |
| LH_Default_PFC_4 | -0.005 | -0.004 | -0.009* | 0.008 | -0.012** |
| LH_Default_PFC_5 | 0.003 | -0.001 | -0.003 | -0.006 | 0.000 |
| LH_Default_PFC_6 | -0.001 | -0.004 | -0.007 | -0.002 | -0.002 |
| LH_Default_PFC_7 | -0.006 | -0.009 | -0.012** | -0.006 | -0.008* |
| LH_Default_Par_1 | 0.007 | 0.025*** | 0.014*** | 0.018*** | 0.019*** |
| LH_Default_Par_2 | 0.002 | 0.001 | -0.009* | -0.001 | -0.002 |
| LH_Default_Temp_1 | 0.011* | 0.008 | 0.004 | -0.001 | 0.013*** |
| LH_Default_Temp_2 | 0.011* | 0.015*** | 0.006 | 0.007 | 0.012** |
| LH_Default_pCunPCC_1 | 0.002 | 0.002 | -0.014*** | 0.001 | 0.001 |
| LH_Default_pCunPCC_2 | 0.002 | 0.009* | -0.01** | 0.006 | 0.002 |
| LH_DorsAttn_FEF_1 | -0.004 | -0.013** | -0.012** | -0.016* | -0.002 |
| LH_DorsAttn_Post_1 | -0.003 | 0.009* | -0.001 | 0.005 | 0.006 |
| LH_DorsAttn_Post_2 | -0.011* | -0.015** | -0.011** | -0.01* | -0.008 |
| LH_DorsAttn_Post_3 | -0.001 | -0.011* | -0.013** | -0.008 | 0.001 |
| LH_DorsAttn_Post_4 | -0.01* | -0.009* | -0.009* | -0.001 | -0.01* |
| LH_DorsAttn_Post_5 | -0.004 | 0.004 | -0.015*** | -0.001 | 0.000 |
| LH_DorsAttn_Post_6 | -0.01 | -0.011* | -0.015*** | -0.009 | -0.007 |
| LH_DorsAttn_PrCv_1 | -0.005 | -0.008 | -0.005 | -0.005 | 0.002 |
| LH_Limbic_OFC_1 | -0.002 | 0.012** | -0.005 | 0.025*** | -0.004 |
| LH_Limbic_TempPole_1 | 0.005 | 0.006 | 0.001 | 0.003 | 0.01** |
| LH_Limbic_TempPole_2 | -0.008 | -0.012** | -0.012** | -0.011* | -0.009* |
| LH_SalVentAttn_FrOperIns_1 | 0.002 | -0.013*** | -0.01** | -0.018*** | 0.003 |
| LH_SalVentAttn_FrOperIns_2 | -0.004 | -0.017*** | -0.01** | -0.018*** | -0.006 |
| LH_SalVentAttn_Med_1 | -0.005 | -0.017*** | -0.014*** | -0.016** | -0.011** |

| | | | | | |
|---|---|---|---|---|---|
| LH_SalVentAttn_Med_2 | -0.004 | -0.009 | -0.019*** | -0.011* | -0.008 |
| LH_SalVentAttn_Med_3 | -0.002 | -0.011* | -0.008* | -0.017** | -0.005 |
| LH_SalVentAttn_PFCl_1 | -0.004 | -0.009 | -0.009* | -0.010 | -0.01* |
| LH_SalVentAttn_ParOper_1 | 0.001 | 0.003 | -0.001 | -0.002 | 0.002 |
| LH_SomMot_1 | 0.012* | 0.013*** | -0.002 | 0.004 | 0.006 |
| LH_SomMot_2 | -0.004 | -0.013*** | -0.018*** | -0.009 | -0.009* |
| LH_SomMot_3 | 0 | -0.006 | -0.013** | -0.01* | -0.004 |
| LH_SomMot_4 | -0.011* | -0.015*** | -0.016*** | -0.009 | -0.006 |
| LH_SomMot_5 | -0.007 | -0.014** | -0.01** | -0.014* | -0.009* |
| LH_SomMot_6 | -0.008 | -0.012* | -0.016*** | -0.015* | -0.012** |
| LH_Vis_1 | 0 | -0.009* | -0.01** | -0.01* | 0.002 |
| LH_Vis_2 | 0.007 | 0.019*** | -0.002 | 0.007 | 0.008 |
| LH_Vis_3 | 0.006 | 0.009* | -0.01* | 0.003 | 0.003 |
| LH_Vis_4 | 0.002 | 0.008 | 0.002 | -0.001 | 0.004 |
| LH_Vis_5 | 0.003 | 0.019*** | -0.001 | 0.003 | 0.003 |
| LH_Vis_6 | 0.01* | 0.006 | -0.011** | -0.002 | 0.002 |
| LH_Vis_7 | 0 | 0.014** | 0.007 | 0.008* | 0.010 |
| LH_Vis_8 | 0.003 | 0.008 | -0.001 | 0.005 | 0.008 |
| LH_Vis_9 | 0.005 | 0.006 | -0.02*** | -0.006 | -0.005 |
| RH_Cont_Cing_1 | -0.003 | -0.01* | -0.013*** | -0.01* | -0.013** |
| RH_Cont_PFCl_1 | -0.003 | -0.002 | -0.01** | 0.01* | -0.008* |
| RH_Cont_PFCl_2 | -0.011* | -0.011* | -0.008* | 0.003 | -0.005 |
| RH_Cont_PFCl_3 | -0.006 | -0.006 | -0.011** | -0.002 | -0.011** |
| RH_Cont_PFCl_4 | -0.004 | 0.003 | -0.005 | 0.010 | 0.001 |
| RH_Cont_PFCmp_1 | -0.006 | -0.012** | -0.011** | -0.007 | -0.009* |
| RH_Cont_Par_1 | 0.002 | -0.002 | -0.004 | -0.004 | -0.004 |
| RH_Cont_Par_2 | 0.001 | -0.005 | -0.011** | -0.005 | -0.003 |
| RH_Cont_pCun_1 | -0.005 | 0.003 | -0.018*** | 0.003 | -0.004 |
| RH_Default_PFCdPFCm_1 | -0.001 | 0.000 | -0.007 | 0.004 | -0.002 |
| RH_Default_PFCdPFCm_2 | 0.004 | 0.007 | 0.002 | 0.004 | 0.008 |
| RH_Default_PFCdPFCm_3 | -0.006 | -0.002 | -0.011** | 0.002 | -0.004 |
| RH_Default_PFCv_1 | 0 | 0.004 | -0.003 | 0.016*** | 0.000 |
| RH_Default_PFCv_2 | 0.003 | 0.017*** | 0.012*** | 0.028*** | 0.014*** |
| RH_Default_Par_1 | 0.005 | 0.016*** | 0.003 | 0.014*** | 0.006 |
| RH_Default_Temp_1 | 0.001 | 0.019*** | -0.004 | 0.028*** | 0.001 |
| RH_Default_Temp_2 | 0.01* | 0.026*** | 0.011** | 0.025*** | 0.02*** |
| RH_Default_Temp_3 | 0.013* | 0.031*** | 0.013** | 0.024*** | 0.016*** |
| RH_Default_pCunPCC_1 | 0.004 | 0.009* | -0.012** | 0.008* | 0.005 |

| | | | | | |
|---|---|---|---|---|---|
| RH_Default_pCunPCC_2 | -0.001 | 0.011** | -0.008* | 0.011** | 0.004 |
| RH_DorsAttn_FEF_1 | -0.004 | -0.009 | -0.009* | -0.010 | 0.000 |
| RH_DorsAttn_Post_1 | 0.004 | 0.016*** | 0.003 | 0.013** | 0.014** |
| RH_DorsAttn_Post_2 | -0.004 | -0.006 | -0.005 | 0.002 | -0.004 |
| RH_DorsAttn_Post_3 | -0.004 | -0.013** | -0.011** | -0.010 | -0.006 |
| RH_DorsAttn_Post_4 | -0.001 | -0.007 | -0.012** | -0.005 | 0.001 |
| RH_DorsAttn_Post_5 | -0.005 | -0.001 | -0.014*** | -0.006 | -0.001 |
| RH_DorsAttn_PrCv_1 | -0.002 | 0.008 | 0.003 | 0.018*** | 0.009* |
| RH_Limbic_OFC_1 | -0.001 | 0.016*** | -0.001 | 0.03*** | -0.003 |
| RH_Limbic_TempPole_1 | 0.001 | 0.021*** | 0.003 | 0.029*** | 0.009* |
| RH_SalVentAttn_FrOperIns_1 | -0.006 | -0.014*** | -0.013*** | -0.006 | -0.009* |
| RH_SalVentAttn_Med_1 | -0.004 | -0.01* | -0.019*** | -0.010 | -0.011** |
| RH_SalVentAttn_Med_2 | -0.005 | -0.013** | -0.01** | -0.017** | -0.006 |
| RH_SalVentAttn_TempOccPar_1 | 0.006 | 0.02*** | 0.008 | 0.011** | 0.015*** |
| RH_SalVentAttn_TempOccPar_2 | -0.006 | -0.003 | -0.007 | 0.003 | -0.007 |
| RH_SomMot_1 | 0.006 | 0.006 | -0.012** | 0.004 | -0.004 |
| RH_SomMot_2 | -0.006 | -0.015*** | -0.019*** | -0.007 | -0.011** |
| RH_SomMot_3 | -0.002 | -0.006 | -0.015*** | -0.002 | -0.009* |
| RH_SomMot_4 | -0.007 | -0.007 | -0.016*** | 0.002 | -0.006 |
| RH_SomMot_5 | -0.004 | -0.007 | -0.011** | -0.005 | -0.003 |
| RH_SomMot_6 | -0.01* | -0.015** | -0.01** | -0.014* | -0.008* |
| RH_SomMot_7 | -0.001 | 0.000 | -0.007 | -0.001 | -0.006 |
| RH_SomMot_8 | -0.009 | -0.014** | -0.015*** | -0.017* | -0.011** |
| RH_Vis_1 | -0.002 | 0.001 | -0.011** | 0.004 | 0.002 |
| RH_Vis_2 | 0.008 | 0.022*** | -0.001 | 0.017*** | 0.008 |
| RH_Vis_3 | 0 | 0.018*** | 0.000 | 0.014** | 0.003 |
| RH_Vis_4 | 0.01 | 0.027*** | 0.006 | 0.015*** | 0.008 |
| RH_Vis_5 | 0.012* | 0.023*** | -0.006 | 0.011* | 0.002 |
| RH_Vis_6 | 0.013* | 0.012*** | -0.016*** | 0.006 | -0.001 |
| RH_Vis_7 | 0.005 | 0.01* | 0.003 | 0.01* | 0.010 |
| RH_Vis_8 | 0.007 | 0.018*** | -0.006 | 0.008 | 0.002 |

**Supplementary Table 11.** The EEG-fMRI correlations within each Yeo (2011) network for each frequency band compared between familiar and unfamiliar participants. Values are Fisher's z transformed correlation values. Significance of difference between the groups was assessed with a group assignment permutation test and a false discovery rate correction within a frequency band. Significant differences are denoted *** = $p$ < .001, ** $p$ < .01, * = $p$ < .05.

| | Familiar - Unfamiliar | | | | |
|---|---|---|---|---|---|
| **Network** | **Alpha** | **Beta** | **Delta** | **Gamma** | **Theta** |
| Visual | .005 | .012*** | -.005 | .006 | .004 |
| Dorsal Attention | -.004 | -.005 | -.008* | -.003 | -.000 |
| Ventral Attention | -.003 | -.008 | -.009** | -.009* | -.004 |
| Default Mode | .003 | .007* | -.002 | .008* | .004 |
| Somato-motor | -.004 | -.007 | -.013*** | -.007 | -.007* |
| Fronto-parietal | -.003 | -.006 | .012*** | -.003 | -.007* |
| Limbic | -.001 | .009* | -.003 | .015*** | .001 |

**Supplementary Table 12.** Pearson correlation between brain rhythms in the intersubject correlation over time for the familiar and unfamiliar groups, and the difference between the groups. Significance of the correlation was assessed with a permutation test and a false discovery rate correction across all comparisons. Significant differences are denoted *** = *p* < .001, ** *p* < .01, * = *p* < .05.

| | **Familiar** | **Unfamiliar** | **Familiar - Unfamiliar** |
|---|---|---|---|
| Alpha-Beta | .287*** | .185*** | .102*** |
| Alpha-Gamma | .273** | .150 | .123*** |
| Alpha-Theta | .352*** | .332*** | .020 |
| Alpha-Delta | .114*** | .078*** | .036 |
| Beta-Gamma | .610*** | .442*** | .168*** |
| Beta-Theta | .182*** | .090* | .092*** |
| Beta-Delta | .058 | .085* | -.027 |
| Gamma-Theta | .201*** | .124*** | .077*** |
| Gamma-Delta | .051 | .061 | -.010 |
| Theta-Delta | .405*** | .323*** | .082*** |

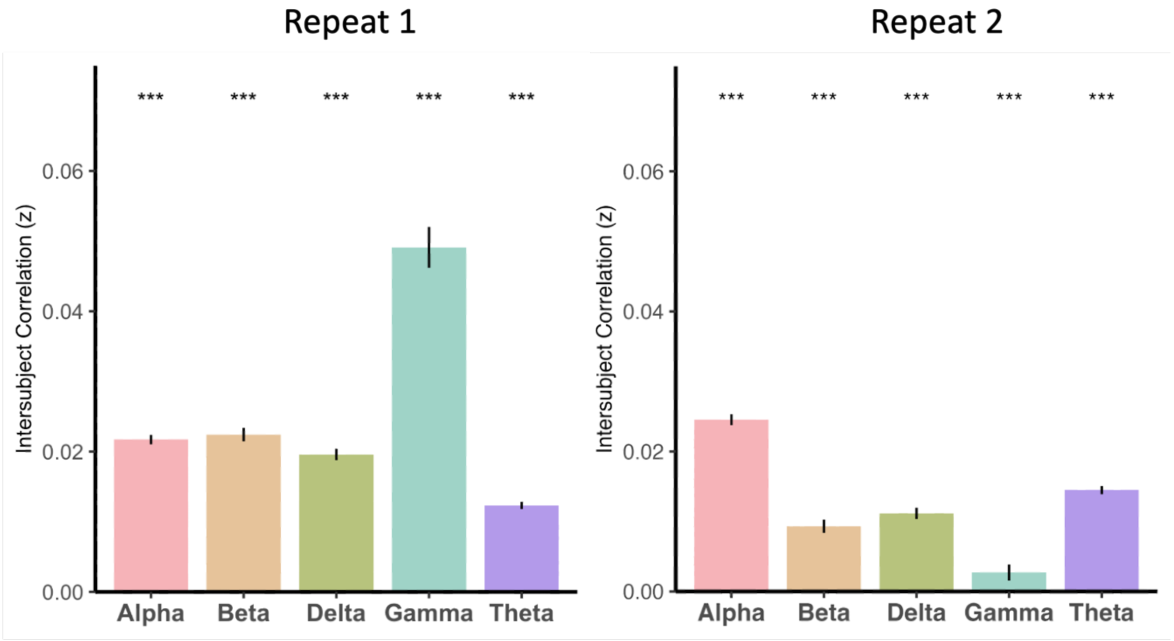


**Supplementary Figure 1.** Intersubject correlations for the first showing of the movie stimulus compared to the second repeat across all participants. ISCs remain significantly above zero in the second repeat, but beta, delta and gamma are significantly attenuated compared to the first viewing.

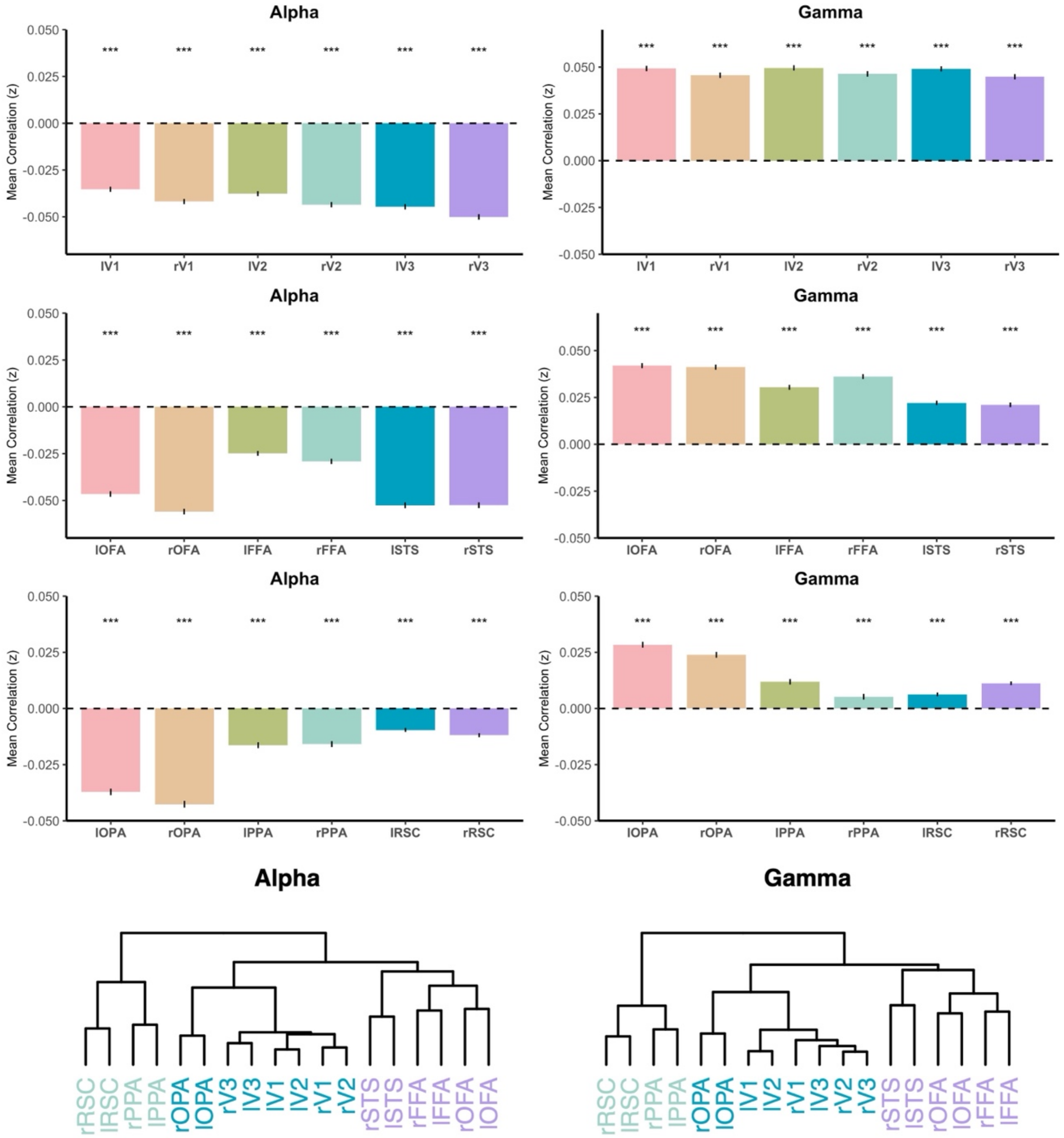


**Supplementary Figure 2.** EEG-fMRI correlations in alpha and gamma in early visual, face- and place-selective cortex. Alpha and gamma show opposite patterns (alpha negatively correlation, gamma positively correlated), but the organisation of these relations is highly similar.

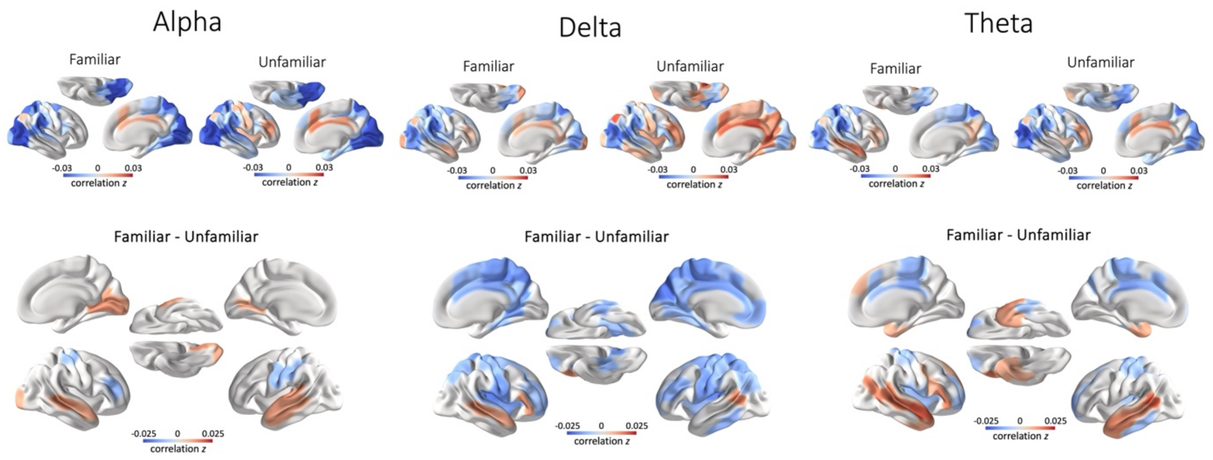


**Supplementary Figure 3.** EEG-fMRI correlations for alpha, delta and theta split by group familiarity. Some global group differences are found in delta, characterised by greater delta-BOLD correlations in unfamiliar participants in non-visual brain regions. Theta demonstrates some group differences, characterised by more positive correlations with BOLD in the temporal lobe of familiar participants.

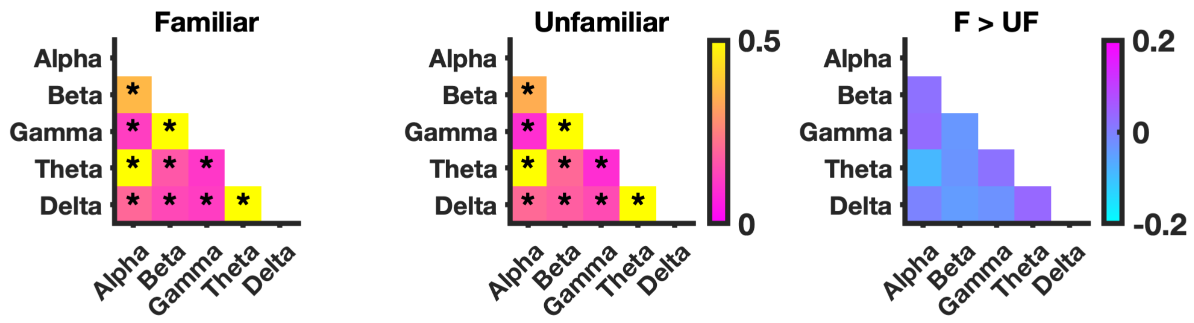


**Supplementary Figure 4.** Within-subject correlations between brain rhythms split during movie watching for familiar and unfamiliar participants. All bands showed significantly greater correlations than the null distribution (corrected for multiple comparisons). No significant differences in correlations between rhythms were found between familiar and unfamiliar participants. This analysis did not reproduce the differences in coupling revealed by the more sensitive spatial ISC analysis (Fig. 3).
